# Genomic and epigenomic adaptation in SP-R210 (Myo18A) isoform-deficient macrophages

**DOI:** 10.1101/2021.04.02.438271

**Authors:** Eric Yau, Yan Chen, Chunhua Song, Jason Webb, Marykate Carillo, Yuka Imamura Ikawasawa, Zhenyuan Tang, Yoshinori Takahashi, Todd M Umstead, Sinisa Dovat, Zissis C. Chroneos

## Abstract

Macrophages play fundamental roles in regulation of inflammatory responses to pathogens, resolution of inflammation and tissue repair, and maintenance of tissue homeostasis. The long (L) and short (S) isoforms of SP-R210/MYO18A, a macrophage receptor for surfactant protein A (SP-A) and C1q, regulate basal and inflammatory macrophage phenotype at multiple gene expression, translational, and subcellular levels in addition to their SP-A and C1q-mediated functions; disruption of L renders macrophages hyper-inflammatory, although the underlying mechanism had previously been unexplored. We asked whether disruption of the L isoform led to the hyper-inflammatory state via alteration of global genomic responses. RNA sequencing analysis of SP-R210_L_(DN) macrophages revealed basal and influenza induced upregulation of genes associated with inflammatory pathways, including TLR, RIG-I, NOD, and cytoplasmic DNA signaling, whereas knockdown of both SP-R210 isoforms (L and S) only resulted in increased RIG-I and NOD signaling. Chromatin immunoprecipitation sequencing (ChIP-seq) analysis showed increased genome-wide deposition of the pioneer transcription factor PU.1 in SP-R210_L_(DN) compared to WT cells. ChIP-seq analysis of histone H3 methylation showed alterations in both repressive (H3K9me3 and H3K27me3) and transcriptionally active (H3K9me3) histone marks. Influenza A virus (IAV) infection, which stimulates an array of cytosolic and TLR-mediated antiviral mechanisms, resulted in differential redistribution between proximal promoter and distal sites and decoupling of PU.1 binding from Toll-like receptor regulated gene promoters in SP-R210_L_(DN) cells. Our findings suggest that SP-R210_L_-deficient macrophages are poised with an open PU.1-primed chromatin conformation to rapidly respond to inflammatory and metabolic stimuli.

## 2 Introduction

Macrophage functions are dynamic, alternating between pro-inflammatory responses to pathogens and stress, and anti-inflammatory responses to alleviate injury to facilitate tissue repair (Sica and Mantovani, 2012, Italiani and Boraschi, 2014, Stout, et al., 2005). Controlling this plasticity is particularly important in the context of alveolar macrophages, the resident immune cell of the lungs (Hussell and Bell, 2014, McQuattie-Pimentel, et al., 2021). As our lungs are exposed to pollutants, pathogens, and particulates on a daily basis that can result in injury, infection, and damage, this fluidity to regulate inflammatory activation state is particularly important for alveolar macrophages to maintain lung homeostasis and avoid overt inflammation (Upham, et al., 1995, Kobzik, et al., 1990). GM-CSF, the transcription factors PU.1 and PPARγ, and the local microenvironment consisting of surfactant proteins and lipids drive alveolar macrophage (AM) development, differentiation, and function (McQuattie-Pimentel, et al., 2021, Shibata, et al., 2001, Baker, et al., 2010, Chroneos, et al., 2010, Guilliams, et al., 2013, Bates, et al., 1997, Schneider, et al., 2014).

Unlike circulating monocyte-derived macrophages, alveolar macrophages are derived from monocyte progenitors in the yolk sac and fetal liver, which then migrate to the lung during fetal development (Guilliams, et al., 2013, Tan and Krasnow, 2016). Early studies depleting alveolar macrophages in mice showed that they play a critical role in pathogen clearance, similar to circulating monocyte-derived macrophages. AMs secrete anti-microbial cytokines such as type I IFNs, phagocytose pathogens, and act as antigen presenting cells to bridge the innate and adaptive immune systems (He, et al., 2017, Kirby, et al., 2009). Depletion of alveolar macrophages also results in overt inflammatory responses in mice exposed to sensitizing proteins, in particular through secretion of cytokines like TGFβ, IL-10, and prostaglandins (Roth and Golub, 1993). Alveolar macrophages are critical for host survival and resolution of influenza infection (Schneider, et al., 2014, Purnama, et al., 2014, Halstead, et al., 2018, Halstead and Chroneos, 2015). AMs are thus critical in maintaining the anti-inflammatory state of the lung under normal conditions, and to resolve inflammation that occurs during pulmonary infections caused not only by the pathogen itself, but also by the host inflammatory response.

The unique microenvironment of the lung alveolus consists of surfactant proteins that work in concert with alveolar macrophages to regulate immune balance, host defense, and tissue homeostasis (Chroneos, et al., 2010, Casals, et al., 2019, Minutti, et al., 2017). Surfactant proteins are part of the lipoprotein surfactant complex that reduces alveolar surface tension to maintain alveolar gas-exchange, alveolar stability, and alveolar recruitment during breathing (Canadas, et al., 2020, Autilio and Perez-Gil, 2019). There are two hydrophobic (SP-B, SP-C) and two hydrophilic lipid-binding (SP-A, SP-D) surfactant proteins, all having amino-terminal domains that are important for oligomerization into lipid-dependent and lipid-independent supramolecular structures. SP-B and SP-C are critical for the formation of the surface tension lowering surfactant monolayer at the air-liquid interface, whereas SP-A and SP-D modulate ultrastructural organization of the surfactant phospholipid sub-phase. Within this milieu, the surfactant proteins enhance pathogen clearance and coordinate immune and metabolic functions of alveolar macrophages, alveolar epithelial cells, and their cross-talk with innate and adaptive immune cells. Under basal conditions, SP-A maintains alveolar macrophages at an anti-inflammatory state by a number of mechanisms that include increasing expression of the transcription factor IRAK-M (Nguyen, et al., 2012), global effects on macrophage proteome composition enriched in anti-inflammatory pathways (Phelps, et al., 2013, Phelps, et al., 2011), scavenging of pro-inflammatory mediators (Minutti, et al., 2016, Francisco, et al., 2020), suppression of NFκB activation (Younis, et al., 2020, Moulakakis, et al., 2016, Moulakakis and Stamme, 2009, Wu, et al., 2004), and modulating trafficking of innate receptors (Henning, et al., 2008, Gil, et al., 2009).

SP-A modulates the optimal relative expression of the long and short isoforms of the SP-A receptor, SP-R210_L_ and SP-R210_s_, on alveolar macrophages (Nguyen, et al., 2012, Yang, et al., 2015). SP-R210 tailors SP-A-mediated phagocytosis of SP-A opsonized pathogens and pathogen-dependent inflammatory responses by macrophages (Weikert, et al., 2000, Weikert, et al., 1997, Minutti, et al., 2017). These isoforms are encoded by alternatively spliced mRNAs of the *Myol8A* gene (Mori, et al., 2003, Yang, et al., 2005, Szeliga, et al., 2005), which generate tissue and cell-type specific isoforms on the cell surface and subcellular organelles (Mori, et al., 2003, Yang, et al., 2005, Szeliga, et al., 2005, Taft and Latham, 2020, Lee, et al., 2014, Ng, et al., 2013, Horsthemke, et al., 2019, Cross, et al., 2004). MYO18A isoforms have so far been classified into three broad groups, MYO18Aα, MYO18Aβ, and MYO18Aγ, which are functionally diverse, catalytically inactive members of the myosin superfamily. SP-R210_L_ (aka CD245α) and SP-R210_s_ (aka CD245β) are members of the MYO18Aα and MYO18Aβ groups on the surface of macrophages and other immune cells (Yang, et al., 2005, De Masson, et al., 2016, Samten, et al., 2008), respectively. SP-R2120_L_ is induced during terminal macrophage differentiation(Mori, et al., 2003). SP-R210/MYO18A isoforms are differentially expressed with different relative abundance in myelomonocytic lineage cells (Mori, et al., 2003, Cross, et al., 2004, Samten, et al., 2008, Chroneos and Shepherd, 1995), although the role of each isoform and impact of SP-R210 isoform stoichiometry in macrophage development and function is not understood at the molecular, cellular levels, and organismal levels. Bone marrow-derived macrophages, monocytes, and immature myelomonocytic cells only express SP-R210_s_, whereas peritoneal and alveolar macrophages express both isoforms (Yang, et al., 2005, Cross, et al., 2004, Samten, et al., 2008). Previous studies demonstrated that SP-R210 regulates extrinsic ligand-dependent and intrinsic ligand-independent macrophage functions (Yang, et al., 2015, Weikert, et al., 2000, Weikert, et al., 1997, Minutti, et al., 2017, Mori, et al., 2003, Yang, et al., 2005, Cross, et al., 2004, Samten, et al., 2008, Jean Beltran, et al., 2016, Sever-Chroneos, et al., 2011, Borron, et al., 1998, Chroneos, et al., 1996, Lopez-Sanchez, et al., 2010). Ligand-dependent functions include SP-A-mediated bacterial phagocytosis coupled to production of reactive nitrogen species, secretion of TNFα and increased macrophage responsiveness to IFNγ and IL-4. These functions facilitate killing and eradication of bacterial and parasitic pathogens by alveolar macrophages (Weikert, et al., 2000, Weikert, et al., 1997, Minutti, et al., 2017, Sever-Chroneos, et al., 2011, Stamme, et al., 2000). In the context of an immune memory response, however, the interaction of SP-A with SP-R210 may limit excessive activation of immune system-mediated inflammation (Samten, et al., 2008, Borron, et al., 1998). SP-R210 may exacerbate injurious inflammation by peritoneal macrophages in the presence of other ligands such as C1q (Minutti, et al., 2017), which is more abundant systemically, and innate immune cell cytotoxicity by SP-A or other ligands in different tissues (De Masson, et al., 2016).

Studies in SP-R210_L_-deficient macrophages revealed that depletion of the L isoform resulted in broad baseline alterations in expression, trafficking and cell-surface localization of innate receptors, and hyper-responsiveness to inflammatory stimuli (Yang, et al., 2015), suggesting global effects on macrophage phenotype and function. However, the role that SP-R210 isoforms play in macrophage function are not well-defined. Therefore, we asked whether differences in SP-R210 isoform expression patterns shape the basal macrophage phenotype and responses to infection at genomic, transcriptomic, and epigenomic levels. We found that alteration of SP-R210 isoform expression resulted in distinct gene expression patterns that reflected changes in epigenetic remodeling and chromatin accessibility in SP-R210_L_-depleted macrophages. Additionally, using influenza as a model of infection, our findings support the notion that SP-R210 isoforms coordinate nuclear responses to infection in macrophages.

## 3 Materials and Methods

### 3.1 Cell lines and Cell Culture

SP-R210_L_-deficient RAW 264.7 macrophages (SP-R210_L_(DN)) were generated and characterized as described previously by stable transfection of a pTriex-2 vector expressing the carboxy-terminal domain of SP-R210 (SP-R210_L_(DN)) cells (Yang, et al., 2015, Yang, et al., 2005, Sever-Chroneos, et al., 2011). Most experiments utilized the SP-R210_L_(DN) clone DN2 (Yang, et al., 2015, Yang, et al., 2005, Sever-Chroneos, et al., 2011) unless otherwise noted in Figure legend. Control cells were transfected with empty vector. The deletion of both SP-R210 isoforms was achieved using CRISPR/Cas9. Guide RNA sequences targeting exons 5, 6, 12, and 14 were designed using crispr.mit.edu and gRNA oligonucleotides were ligated into the LRG hU6-sgRNA-EFS-GFP-P2A vector. RAW 264.7 macrophages were transduced with a Cas9 containing lentivirus and selected using blasticidin to generate a stable Cas9 expressing cell line. Stable Cas9 RAW264.7 cells were transfected with a pooled library of ligated gRNA vectors. Transfected GFP+ cells were isolated by flow activated cell sorting into 96-well plates to culture individual clones. SP-R210-deficient clones lacking both isoforms, designated, SP-R210(KO), were identified by Western blot analysis following sorting (Supplemental Figure 1), and a single clone retaining deletion of both isoforms after subculture was selected for further studies. Cells were maintained in DMEM culture media (DMEM with 4.5 g/L glucose, L-glutamine, and sodium pyruvate supplemented with 10% heat-inactivated fetal bovine serum (FBS) and 1% penicillin/streptomycin) at 37°C and 5% CO_2_ in 10-cm dishes. Cells were sub-cultured in 10-cm dishes x 5 at a density of 1 x 10^7^ cells/dish for ChIP-seq experiments, sub-cultured in 6-well plates at a density of 1 x 10^6^ cells/well for RNA isolations (subjected to RNA-seq experiments), or 24-well plates at 2 x 10^5^ cells/well for flow cytometry and other assays.

### 3.2 Virus Preparation

The mouse adapted influenza virus strain A/Puerto Rico/8/34 (PR8) influenza was propagated in the allantoic fluid of embryonated chicken eggs (Sever-Chroneos, et al., 2011). Briefly, 10^5^ fluorescent focus units (FFC) in PBS with 1% Penicillin/Streptomycin/Fungizone was injected into the amniotic sac of 10 day old embryonated chicken eggs. Infected eggs were incubated at 37°C for 56 hours. Eggs were then removed and placed at 4°C for 12 hours. The allantoic fluid was collected and spun down at 131,000g for 40 minutes at 4°C. The virus pellet was reconstituted in PBS and layered over a 30%/35%/50%/60% sucrose gradient and spun at 168,000g for 1 hour and 15 minutes and the virus containing layer between 50% and 35% was collected and dialyzed against PBS at 4°C overnight. The viral titer was determined by fluorescent plaque assays using Madine-Darby Canine Kidney (MDCK) epithelial cells (ATCC Cat #CRL-2936) were plated in 96 well plates at a density of 3×10^4^ serial dilutions of purified and dialyzed virus was overlayed on the cells. Cells were incubated at 37°C with virus for 2 hours, at which point virus-containing media was replaced with virus-free media and cells were incubated for another 6 hours. Cells were then fixed with acetone, and stained with an Influenza A NP antibody (Sigma Aldrich Cat#MAB8251, 1:100 in PBS) for 30 minutes at 4°C and subsequently labeled with Rhodamine conjugated anti-mouse IgG (Jackson ImmunoResearch, Cat#115-026-062, 1:100 in PBS). Fluorescently labeled nuclei were counted using a Nikon Eclipse TE2000-U at 20x magnification.

### 3.3 Influenza Infection and Assessment of Infection

Cells plated for infection were seeded at a density of 1×10^7^ (ChIP-seq) or 2E5 per well for 12 hours prior to infection. Cells were washed with PBS twice and PR8 virus was added at MOI 1 (ChIP-seq) or MOI 4 (cytokine elaboration studies, influenza infection studies) in infection media (1:1 ratio DMEM w/o serum to PBS) as determined by cell seeding density. Cells with infection media were incubated at 37°C for two hours. Infection media were removed and replaced with DMEM culture media. Infection was allowed to progress for another 10 hours or until the desired incubation time was reached. Influenza infection was assessed via viral protein production by western blot (methods outlined in 2.4), viral NP qPCR, and viral NP imaging. Supernatant and cell pellets from infected samples were collected 12 hours post infection. qPCR was performed in accordance to Fino et al (Fino, et al., 2017). RNA was prepared using the RNA-bee (Tel-Test, Inc. Cat #CS-501B) protocol for cell supernatant and cell pellet. cDNA was synthesized with the High Capacity cDNA Reverse Transcription Kit (Invitrogen Cat#4368814) using the IAV MP primer 5’ TCT AAC CGA GGT CGA AAC GTA 3’ for IAV following the manufacturer’s protocol. The cDNA was diluted five-fold prior and PCR amplified using the TaqMan Fast Universal PCR Master Mix (ThermoFisher Cat#4305719). The PR8 M1 gene was amplified using the following primers: sense: 5’-AAG ACC AAT CCT GTC ACC TCTG A-3’ and antisense: 5’-CAA AGC GTC TAC GCT GCA GTC-3’, 900 nM each and the 200 nM probe sequence: 5’-/56-FAM/ TTT GTG TTC ACG CTC ACC GT/36-TAMSp/-3’. Immunofluorescence of lAV-infected cells was performed on PR8 infected cells PR8 at MOI10 after 15 and 30 minutes. Infected cells were fixed using 3.7% formaldehyde for 25 minutes. Fixed cells were blocked with 10% donkey serum and 3% BSA in 0.2% Tween 20 in PBS. Blocked cells were probed using Rab5 (Cell Signaling Technology Cat#3547S, 1:200) and NP (BioRad Cat#MCA400, 1:1000) overnight at 4°C, and labeled using AF488 conjugated anti-mouse IgG (Jackson ImmunoResearch Cat#715-545-150, 1:500) and AF594 conjugated antirabbit IgG (Jackson ImmunoResearch Cat#711-585-152, 1:500) for 2 hours at room temperature. Cells were acquired using Nikon Eclipse T*i* confocal microscope and Nikon Elements 4.30.01 software.

### 3.4 Western blot analysis

Samples were harvested using 0.25% Trypsin-EDTA (Corning Cat#25-053-Cl); lifted cells were centrifuged at 15,000g for 5 minutes at 4°C (Eppendorf 5430R). Cell pellets were placed at −20°C overnight with 60 μL SDS Lysis buffer (1% SDS, 50mM Tris-HCL pH 8.1, 10mM EDTA pH 8.0) with 1x Protease/Phosphatase Inhibitor Cocktail (Cell Signaling Cat#5872S). Samples were thawed and sonicated at 50% A for 10 seconds in 2 second ON/OFF intervals. Sonicated samples were centrifuged at 15,000g for 5 minutes at 22°C. Supernatant was collected; 4x LDS sample buffer (Invitrogen Cat#NP007) was added to a final concentration of 1x.

For gel electrophoresis, samples were diluted to 1X SDS-PAGE sample buffer, boiled at 90°C for 2 minutes, and centrifuged for 2 minutes at 21,000g. Equal volumes of protein were via in 4-12% BisTris gels (Invitrogen, Cat#NP0321BOX, NP0349BOX) for reducing gels and 4-12% Bis0Tris NativePAGE gels (Invitrogen, Cat #BN1002BOX) for native, nonreducing gels and transferred using eBlot L1 Transfer system (GenScript). Blots were blocked in 5% Bovine Serum Albumin (BSA) in 0.1% Tween 20 in Tris-buffered saline (0.1% TBST). Blots were probed at 4°C overnight with MYO18A (Proteintech Cat#14611-1-AP, 1:1000), p-IRF3 Ser396 (Cell Signaling Technology Cat#29047, 1:1000), IRF3 (Cell Signaling Technology Cat#4302, 1:1000), p-IRF7 Ser437 (Cell Signaling Technology Cat#24129, 1:1000), IRF7 (Aviva Systems Biology Cat#OAAN00009), p-p65 Ser276 (Thermo Fisher Cat#PA5-37718, 1:1000), p-p65 Ser536 (Cell Signaling Technology Cat#3033, 1:1000), NFκB p65 (Cell Signaling Technology Cat#8242, 1:1000), STING (Cell Signaling Technology Cat#50494, 1:1000), or GAPDH (Thermo Fisher Cat#PA1-16777, 1:5000) in 0.1% BSA in 0.1% TBST and subsequently imaged using HRP-conjugated anti-rabbit antibodies (Bio-Rad Cat#172-1019, 1:10000) and Western Lightning Plus-ECL (PerkinElmer Cat#El103001EA) on a BioRad Chemidoc imager. Images were adjusted and band densitometry was determined using ImageLab 5.2.1. PU.1 (Invitrogen Cat #MA5-15064, 1:500-1000), NS1 (Invitrogen Cat #PIPA532243, 1:1000), NP (BioRad Cat#MCA400, 1:1000), and anti-β-actin (Sigma Aldrich Cat #A2228-100UL, 1:20000) blots were incubated with the relevant primary antibodies at 4°C overnight and imaged using with IRDye 680RD anti-Mouse IgG (LI-COR Cat#926-68070, 1:15000) and IRDye 800 CW anti-Rabbit IgG (LI-COR Cat#926-32211, 1:10000). Blots were imaged using LI-COR Odyssey CLx and Image Studio 4.0. Band densitometry was acquired and images were adjusted using Image Studio Lite 5.2.5.

### 3.5 Cytokine analysis

Cytokines were assessed from clarified supernatant using R&D Quantikine ELISA kits for TNFα (R&D Systems, Cat#MTA00B) and DuoSet ELISA kits for IFNβ (R&D Systems, Cat#DY8234-05). 500 μL of supernatant was collected from cell culture experiments, and spun at 15,000g for 10 minutes to remove debris. Clarified supernatants were aliquoted and frozen before utilizing in ELISA experiments. ELISAs were performed following manufacture protocol without diluting supernatant samples.

### 3.6 Flow cytometry

Control, SP-R210_L_(DN), and SP-R210(KO) RAW 264.7 cells were detached using non-enzymatic cell dissociation medium (Sigma-Aldrich Cat#C1544-100ML) and washed in PBS containing 2% fetal bovine serum (FBS). Cells were washed by centrifugation and discarding of supernatant, then blocked with mouse Fc block (BD Biosciences Cat#553142) in PBS at a concentration of 12.5 μg/mL and 2% FBS for 10 minutes at room temperature. After blocking, cells were washed and stained with recommended concentrations of monoclonal antibodies for 30 min at 4°C. Staining was divided into two panels; Cells were washed and placed into HBSS with 2% FBS and 0.02% sodium azide until assessment via BD LSRII flow cytometer. A minimum 30,000 events were collected and data were analyzed via FlowJo 9.8.8. Antibodies used are as follows; CD204 (Bio-Rad Cat#MCA1322A488T, 2F8); MHC II (eBioscience Cat#86-5321-42, M5/114.15.2); CD11b (BioLegend Cat#101242, M1/70); Ly6C (BD Cat#561237, AL-21); TLR2 (eBioscience Cat#12-9021-82, 6C2); F4/80 (eBioscience Cat#25-4801-82, BM8); CD36 (BD Horizon Cat#585933, CRF D-2712); SIRPα (BD Optibuild Cat#742205, P84); CD11c (eBioscience Cat#17-0114-82, N418); TLR4 (eBioscience Cat#12-9041-80, UT41); CD14 (eBioscience Cat#25-0141-82, SA2-8); SiglecF (BD Horizon Cat#562681, E50-2440)

### 3.7 RNA isolation

RNA was isolated from 1 x 10^6^ cells per sample using the RNA-Bee™ (Tel-Test, Inc. Cat#CS-501B) protocol as described in detail previously (Halstead, et al., 2018). Briefly, plated cells were washed with PBS and 0.5 mL of RNA-Bee Isolation Reagent was added to the cells, lysed with repeated pipetting, and transferred to a microcentrifuge tube followed by the addition of 0.1 mL of chloroform was added, mixed by shaking for 15-30 seconds. and centrifuged at 12,000g for 15 minutes at 4°C. The aqueous phase was retained and mixed with 0.5 mL ice-cold isopropanol and placed at −20°C for 3 hours. Samples were then centrifuged at 12,000g for 15 minutes at 4°C and supernatant was discarded. Precipitated RNA was washed with 1 mL ice-cold 75% ethanol twice, and then allowed to air dry for 15-30 minutes at 4°C. RNA was dissolved in 25 μL RNAse, DNAse-free deionized water. RNA concentration and purity were determined by NanoDrop (Thermo Scientific) to confirm an A260:A280 ratio above 1.9. RNA integration number (RIN) was measured using BioAnalyzer (Agilent Technologies) RNA 6000 Nano Kit.

### 3.8 RNA sequencing and analysis

The cDNA libraries were prepared using the NEXTflex™ Illumina Rapid Directional RNA-Seq Library Prep Kit (BioO Scientific) as detailed previously (Halstead, et al., 2018). The libraries were pooled and loaded onto an S1 flow cell on an Illumina NovaSeq 6000 (Illumina) and run for 2X50 cycles at a depth of 25 million reads per sample. De-multiplexed and adapter-trimmed sequencing reads were aligned to the mm10 reference genome using hisat2 (v2.1.0) (Kim, et al., 2015). Abundance for each gene was obtained using featureCounts function available in Rsubread R package (Liao, et al., 2019). The raw count data from 3 independent replicates were analyzed using DESeq2 to obtain differentially expressed genes in uninfected samples, while 2 independent replicates were used for infected experiments (Love, et al., 2014). DESeq2 results were filtered for differentially expressed genes with a p-value of less than 0.05. MGI annotations of immune associated genes were obtained from the gene ontology term 0002376 (http://www.informatics.jax.org/). Differentially expressed immune genes were obtained by filtering the DESeq2 results with p-value <0.05 against the MGI database of immune-associated genes. Genes with p-value less than 0.05 were mapped to KEGG pathways using fgsea (Fast Gene Set Enrichment Analysis) package (Kanehisa and Goto, 2000, Korotkevich, et al., 2019). Ingenuity pathway analysis (IPA, **www.qiagen.com/ingenuity**) was used for upstream pathway analysis of the most highly expressed gene transcripts q<0.2 as measured by Fisher’s exact test.

### 3.9 Chromatin Immunoprecipitation

Five plates each of SP-R210_L_(DN) and WT cells were cultured at a density of 1 x 10^7^ for a total of 5 x 10^7^ cells. Cells were removed from the plates and cross-linked for 10 min with 1% formaldehyde in the growth medium. Chromatin was fragmented using the Bioruptor sonicator (Diagenode) for 30 min (30s pulses, 30s pauses in between for 10 min, run 3 times) to produce fragments ~400nt in size. ChIP assays were performed as reported by Wang et al. (Wang, et al., 2016). Briefly, 25 μL of sheared DNA was aliquoted to run as input DNA. anti-PU.1 antibody (50 μL; Invitrogen Cat #MA5-15064, E.388.3), anti-H3K4me3 (40 μL; Sigma Aldrich Cat#07-449), anti-H3K9me3 (40 μL; Abcam Cat#ab8898), or anti-H3K27me3 (40 μL; Abcam Cat#ab8580) was incubated with 100 μL of washed Goat-anti-rabbit IgG Dynabeads (Invitrogen Cat#11203D) at 4°C overnight in BSA blocking solution (0.5% BSA in PBS). PU.1 antibody-coated Dynabeads (100 μL) were incubated with 500 μL chromatin in TE buffer (with 0.1% deoxycholate, 1% Triton X-100) overnight at 4°C on rotator (Barnstead Labquake Model 4152110). Protein/DNA complexes were washed with RIPA buffer and captured with a Magnetic Particle Concentrator (Invitrogen). DNA-protein crosslink was reversed via incubation at 65°C overnight. Samples were treated with 1 mg/mL proteinase K for 2 hours at 37°. DNA was extracted using phenol and chloroform extraction and precipitated using 100% EtOH. The dried DNA pellet was reconstituted in 50 μL H_2_O treated with 330 μg/mL of RNase A for 2 hours at 37°C and then recovered using the QIAquick PCR Purification kit (QIAGEN Cat#28104). DNA concentration was determined by Qubit (Thermofisher).

### 3.10 ChIP sequencing and analysis

Libraries of ChIP-derived DNA were sequenced via Illumina NGS. Sequences were aligned to the mm10 genome using the mem function of the bwa package. Peaks were identified from the bed files using the callpeaks function of MACS2 (v2.1.0). Peak annotation and identification of overlapping peaks was done via ChIPpeakAnno (v3.14) using the UCSC mm10 annotated genome (Zhu, et al., 2010). Concordant peaks between the two replicates were selected for further analysis using overlappingPeaks function set at a maxgap=50. Genomic distribution of peaks and pathway analysis was performed using ChIPseeker (v1.16), ReactomePA (v1.24), and clusterProfiler (Yu, et al., 2015, Yu and He, 2016).

### 3.11 Statistical Analysis

Statistical comparison of data was performed using GraphPad Prism 7.0d software (San Diego, CA). 2-way ANOVA with paired samples and unpaired comparisons via *t*-test corrected by the Holm-Sidak method were used to assess statistical differences. p-values<0.05 were considered significant.

## 4 Results

### 4.1 SP-R210 isoform deficient macrophages exhibit basal immune activation pathways

Previous work has shown that knockdown of the longer SP-R210_L_ isoform (SP-R210_L_(DN)) alters cell-surface expression of multiple innate receptor and macrophage differentiation markers (Yang, et al., 2015) (Supplemental Figure 1b). We assessed the impact of SP-R210 isoform deficiency on the basal transcriptome to better understand the phenotypic and functional differences of SP-R210 deficient macrophages. Deseq2 was used on RNA-seq from 3 independent replicates of WT and SP-R210_L_(DN) cells to identify differentially expressed genes. This investigation revealed significant differences in the basal transcriptome profile between WT and SP-R210_L_(DN). Selective deletion of the SP-R210_L_ isoform resulted in upregulation of 1273 genes (p<0.05), with *Fads2, Igf1, Runx3, Cxcl10, Ly86, Ifi44, Ifi44l* among the top 20 genes with the largest expression differences (Figure 1a, Supplementary Table 1). Furthermore, upregulated genes such as *Hhex* and *Il2rg* were identified by their extremely low p-value. Conversely, knockdown of the L isoform resulted in downregulation of 1652 genes, with *Dlg5, Maged1, Bco1, Tmem54 Col5a1*, and *Kdm5d* among the top 20 most downregulated genes (Figure 1a, Supplemental Table 1). Other downregulated genes that were identified by noticeably low p-values included *Ccl6, Lyz1, Myo6, Stard10, and Zcchc24*. Immune associated genes were then identified within the differentially expressed genes by filtering by the gene ontology term GO: 0002376. This filtering identified most of the top upregulated immune associated genes in SP-R210_L_(DN) cells, such as *Tnfsf8, Ifi44, Ifi44l, Ly86, Runx3, Igf1* (Figure 1a (red dots), Figure 1b). There were 250 and 211 immune-associated genes that were upregulated and downregulated in SP-R210_L_(DN) cells compared to WT cells, respectively.

**Figure 1.**
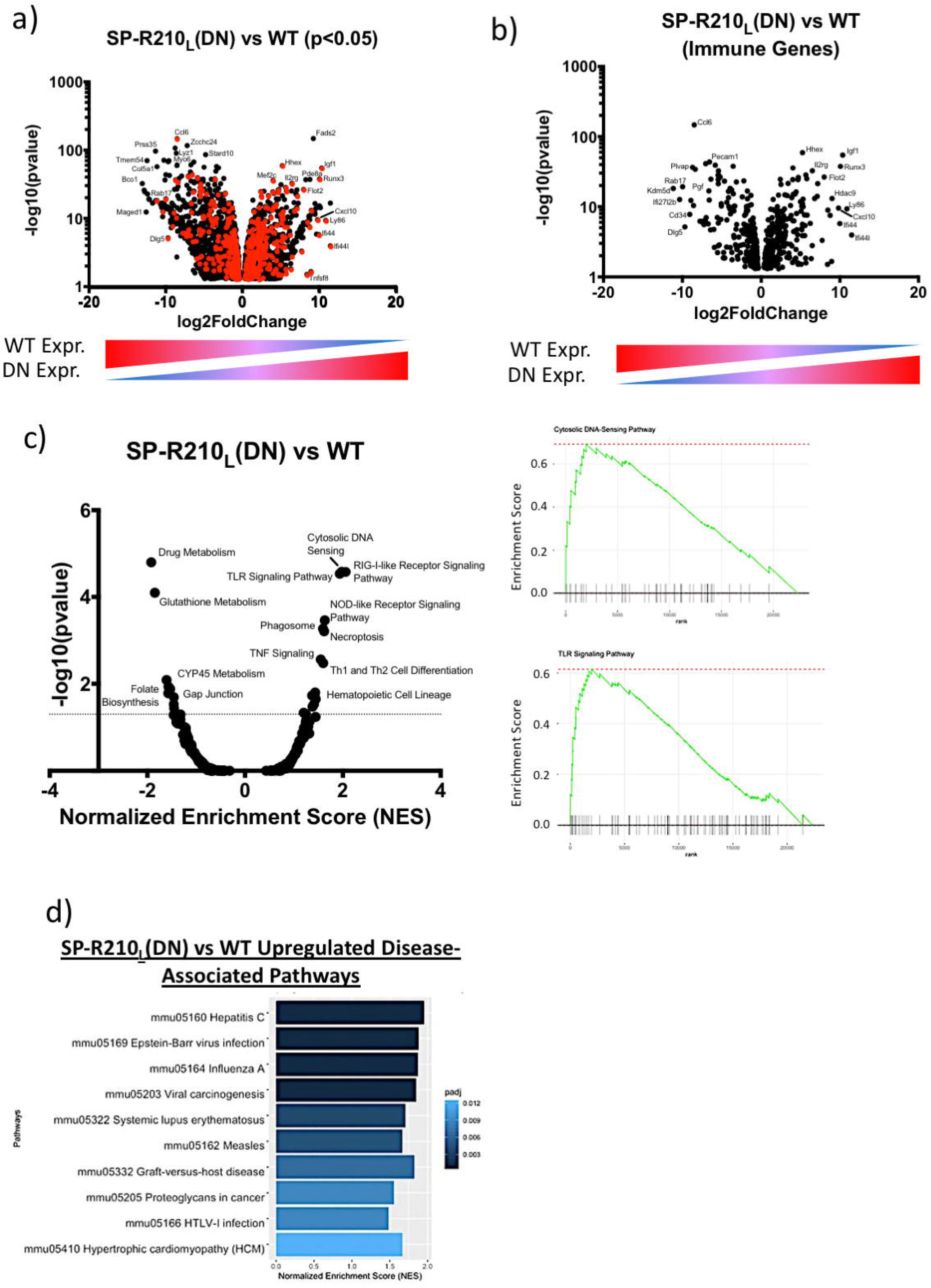
Differentially Expressed genes are associated with upregulation of Innate Immune Sensing Pathways. **(a)** Cells were cultured overnight a 2E5 cells/well and removed using Cell Dissociation Media. Cells were washed, blocked with BD Mouse Fc Block, and stained with fluorescent antibodies against specific cell surface markers. Stained cells were analyzed using a LSR II flow cytometer with compensation and gating analysis performed on FlowJo v 9.9.5. Data plotted is mean of mean fluorescence intensity ± S.E. (n=3), **, adjusted p-value <0.005 compared to WT; ***, adjusted p-value <0.0005 compared to WT. **(b, c)** RNA isolated from WT and SP-R210_L_(DN) cells cultured overnight was sequenced and aligned to the mm10 database using hisat2 with read counts obtained using featurecounts. Count data was then compared between genotypes with three replciates per cell type using deseq2. Differently expressed genes between SP-R210_L_(DN) and WT cells **(b**were filtered by p-value<0.05. These gene sets were then filtered using the MGI Immune Genes database to elucidate differentially expressed immune genes (**c** – SP-R210_L_(DN) vs WT) The genes were also labeled in red in **(b).** Differentially expressed RNA genes were mapped to KEGG pathways using the fgsea R package. **(d)** Upregulated pathways were compared between SP-R210_L_ and WT cells. Enrichment plots for Cytosolic DNA Sensing Pathway and TLR Signaling Pathway were included to show enrichment in genes associated with these gene sets. **(e)** The enrichment scores for the 10 disease associated pathways with the lowest p-values were plotted for SP-R210_L_(DN) vs WT cells.

Differentially expressed genes were then analyzed for enrichment in KEGG signaling/metabolism pathways using the fgsea R package. This analysis revealed induction of cytosolic innate recognition pathways for bacteria, RNA and DNA viruses (NOD, RIG-I, and STING, respectively) in SP-R210_L_(DN) cells as well as cell-surface and endosomal toll-like receptors (Figure 1c, Supplemental Figure 2). Furthermore, several innate sensing pathways (i.e. TLR sensing, NOD sensing) were upregulated in SP-R210_L_(DN) cells, which appears to be skewed towards an anti-viral response due to upregulation of cytosolic DNA sensing pathways and RIG-I. Accordingly, the disease associated pathways most upregulated at baseline in SP-R210_L_(DN) cells were anti-viral response pathways, including responses to Hepatitis C, Epstein-Barr Virus and Influenza A (Figure 1d).

To address the impact of both isoforms, a SP-R210(KO) RAW264.7 macrophage line was generated by CRISPR-Cas9-mediated deletion of both isoforms and expansion of a single clone selected by fluorescence-activated cell sorting (FACS) and deletion confirmed by western blot (Supplemental Figure 1a). Phenotypic analysis by flow cytometry discerned three phenotypes of CD36^+^CD11b^+^F4/80^+^CD14^+^CD204(SR-A)^+^CD284(TLR4)^+^, CD36^high^CD11b^high^F4/80^low^CD14^high^CD204(SR-A)^high^CD284(TLR4)^low^, and CD36^low^CD11b^high^F4/80^+^CD14^++^CD204(SR-A)^++^CD284(TLR4)^++^ for WT, SP-R210_L_(DN), and SP-R210(KO) cells (Supplemental Figure 1b), respectively. TLR2, SIRPα CD11c, Ly6C, and MHC-II were also highly induced in SP-R210_L_(DN) cells while only moderately elevated in SP-R210(KO) cells compared to WT (Supplemental Figure 1b).

Comparative analysis of RNAseq data revealed that disruption of L and both L and S isoforms results in widespread transcriptome adaptation affecting both immune and nonimmune genes. 804 genes were upregulated and 778 downregulated in SP-R210(KO) cells compared to WT. *Tnfsf4, Plch1, Amer3, and Cxcl10* were among the top 20 upregulated genes in the SP-R210(KO) cells, while *Maged1, Zfhx4, Cldn11, and Csf1* were among the 20 most downregulated genes (Supplemental Figure 1c, Supplemental Table 1). Compared to SP-R210_L_(DN) cells, 1936 genes were upregulated while 1525 genes were downregulated in SP-R210(KO)cells compared to SP-R210_L_(DN) cells. Upregulated genes included *Bco1, Tmem54, Ifi27l2b*, and *Kdm5d*, while *Ly86, Igf1, Fads2, Runx3, Ifi44, Csf1, Ifi44l*, and CD86 were among the top 20 downregulated genes (Supplemental Figure 1d, Supplemental Table 1). *Ly86, Ifi44, Ifi44l, Runx3, and Fads2* were all among the top 20 genes that were downregulated in both WT and SP-R210(KO) cells compared to SP-R210_L_(DN) cells. Furthermore, 224 transcripts common to both SP-R210_L_(DN) and SP-R210(KO) were downregulated while 596 genes were upregulated in both cell types compared to WT cells (Supplemental Figure 1g) and of these, 171 upregulated and 101 downregulated transcripts are from immune-associated genes. Among the top 20 upregulated genes in SP-R210(KO) cells, only *Tnfsf4, Itga2, Ifi27l2b*, and *Cxcl10* were immune-associated compared to WT cells (Supplemental Figure 1c, red dots). Tnsfsf4 and *Ifi27l2b* remained elevated in SP-R210(KO) cells to SP-R210_L_(DN) cells (Supplemental Figure 1d, red dots). KEGG pathway analysis showed that only RIG-I and NOD-like receptor signaling pathways remained elevated SP-R210(KO) compared to WT cells-(Supplemental Figure 1e, 1f), whereas nucleic acid sensing pathways were not affected. Taken together, these findings indicate basal macrophage activation readiness at the transcriptional level depends on differential expression of SP-R210 isoforms.

### 4.2 Depletion of SP-R210_L_ alters genome wide binding of PU.1

To further understand the impact of L depletion, chromatin immunoprecipitation and sequencing (ChIP-seq) was used to determine the genome-wide distribution of PU.1 binding in WT and SP-R210_L_(DN) cells. PU.1, a pioneer transcription factor, plays a key role in macrophage function and lineage through its interaction with a constellation of transcription factors at distal gene enhancer and proximal promoter elements, allowing it to prime expression of various macrophage and immune associated genes (Shibata, et al., 2001, Glass and Natoli, 2016, Hoogenkamp, et al., 2007, Imperato, et al., 2015, Petrovick, et al., 1998, Schmidt, et al., 2016, Berclaz, et al., 2007). ChIP-seq was performed on two independent samples for each cell type; concordant peaks were identified between both replicates and were used for further analysis. Comparative analysis of ChIP-seq peaks showed that the number of PU.1 binding sites increased in SP-R210_L_(DN) cells compared to WT (Figure 2a). Furthermore, ChIPseeker analysis revealed that SP-R210_L_(DN) cells had a greater distribution of PU.1 binding in introns than in WT cells (34.64 % vs 33.57%, respectively (Figure 2b). Conversely, binding in promoter regions and regions downstream of gene transcriptional start sites decreased in SP-R210_L_(DN) compared to WT cells (24.94 vs 26.11%, respectively) (Figure 2b). Differences in PU.1 binding were not due to differences in PU.1 expression levels, as demonstrated by Western blot analysis (Supplemental Figure 2a, 2b). These results indicate that knockdown of SP-R210_L_ shifts the distribution of PU.1 binding sites between intergenic and proximal gene promoter sites.

**Figure 2.**
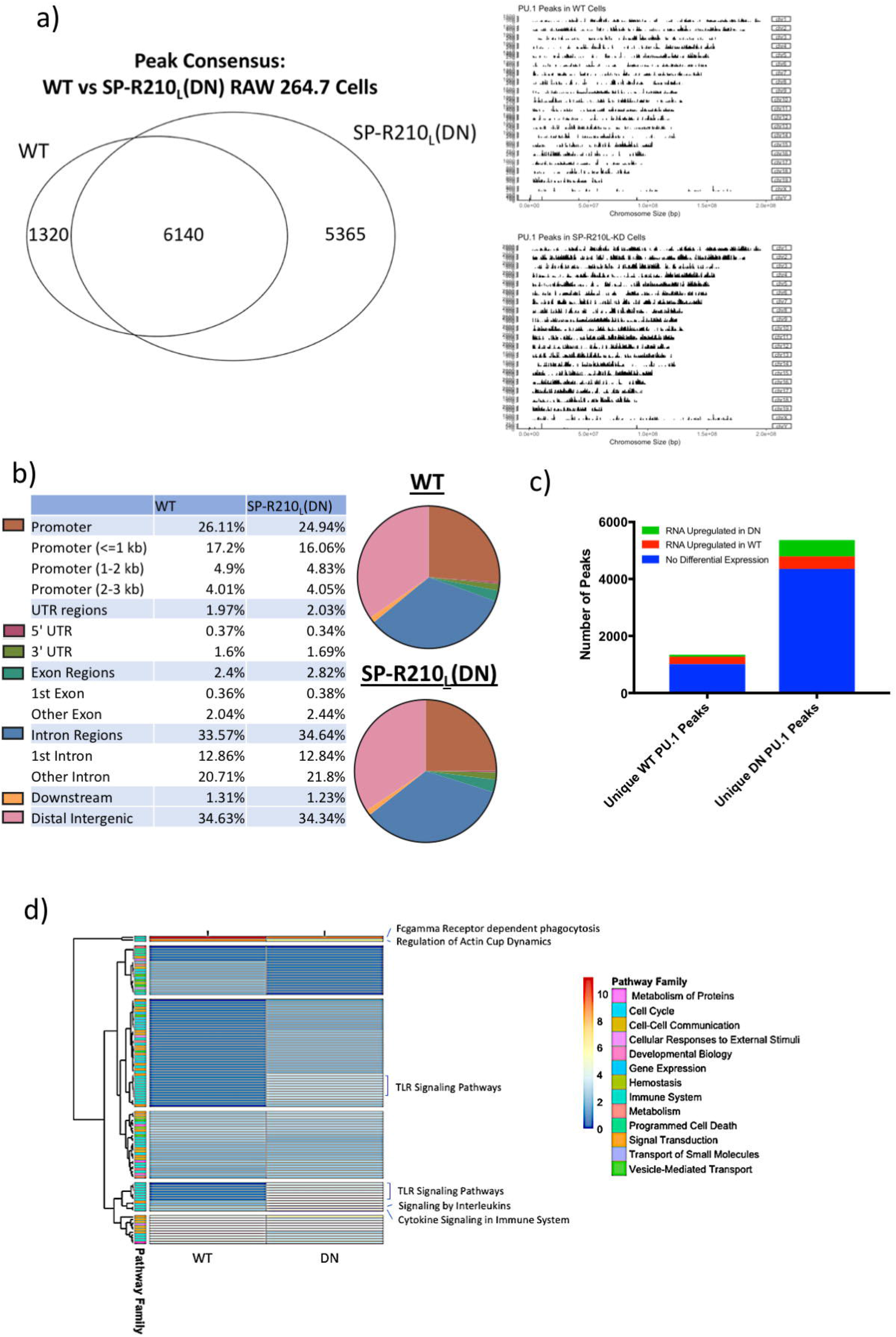
PU.1 binding across the genome is altered with SP-R210_L_ depletion. ChIP was used to precipitate PU.1-bound genomic regions, then sequenced and aligned to the mm10 genome. Concordant peaks between two experimental replicates determined using overlappingpeaks function of ChIPPeakAnno were used for further analysis. **(a)** Peaks between WT and SP-R210_L_(DN) cells were compared using ChIPPeakAnno to identify the concordance in peaks between the two genotypes, showing 6140 peaks consistent with both genotypes, and 5365 peaks unique to SP-R210_L_(DN) cells. The peak distribution across genome was increased in SP-R210_L_(DN) cells **(a)**. **(b)** Using Chipseeker, the identified peaks were associated with genomic features for WT and SP-R210_L_(DN) cells showed decrease PU.1 binding in promoter regions, but with slightly increased binding in 3’ UTR, Exon, and Intron regions. **(c)** Association between unique ChIP peaks in each cell type and RNA expression. Of the 5365 PU.1 peaks unique to SP-R210_L_(DN) cells, 437 peaks were associated with RNA transcripts upregulated in WT cells, while 573 peaks were associated with RNA transcripts upregulated in SP-R210_L_(DN) cells. Of 1011 PU.1 peaks unique to WT cells, 260 peaks were associated with genes with upregulated RNA transcripts in WT cells, and 49 peaks associated with upregulated in SP-R210_L_(DN) cells. **(d)** PU.1 associated genes were mapped to Reactome pathways using ReactomePA. Pathway enrichment scores and p-values for WT and SP-R210_L_(DN) cells were plotted in a heat map; each pathway was associated to larger Reactome pathway families.

The PU.1 peaks were annotated using ChIPseeker, to associate the peaks to specific genes. Of the 1320 peaks unique to WT cells, 260 of the peaks were associated with transcripts downregulated in SP-R210_L_(DN) cells, while 49 peaks were associated with upregulated transcripts in SP-R210_L_(DN) cells. Of the 5365 peaks unique to SP-R210_L_(DN) cells, 573 were associated with upregulated transcripts in the knockdown cells, while 437 were associated with downregulated transcripts in the knockdown cells (Supplemental Table 2a, Supplemental Figure 3). Additionally, of the 1273 upregulated genes in SP-R210_L_(DN) cells, 363 had PU.1 peaks that were found exclusively in SP-R210_L_(DN) cells, with 44 genes having PU.1 peaks exclusive to WT cells. Within the 1652 downregulated genes, 226 genes had PU.1 peaks unique to WT cells, while 308 genes had PU.1 peaks unique to the SP-R210_L_(DN) cells (Figure 2c). Of note, upregulated SP-R210_L_(DN) genes that had PU.1 peaks unique to these cells include several immune associated genes, such as *Ly86, Csf3r, Pde8a*, and *Igf1*, as well as epigenetic regulators such as *Hdac9* (Supplemental Table 2a). On the other hand, downregulated SP-R210_L_(DN) genes that associated with unique PU.1 peaks in the knockdown cells include genes nvolved in signaling, such as *Marcks, Rgs9, Fgr* and *Plekha6* (Supplemental Table 2b).

We then used the ReactomePA package to map gene regions immunoprecipitated by PU.1 to Reactome pathways and clusterProfiler to determine pathway clustering. This analysis showed enrichment in pathways involved in macrophage function in both WT and SP-R210_L_(DN) cells. Pathways regulating Fc gamma receptor dependent phagocytosis, genes regulating actin dynamics for phagocytic cup formation, as well as other immune associated pathways, such as IL-3,5 and GM-CSF signaling, and CD28 co-stimulation pathways (Supplemental Figure 4). SP-R210_L_(DN) cells, however, showed an increased representation of genes involved in TLR pathways (Figure 2d, Supplemental Figure 5), consistent with the RNA-seq data showing differential expression of transcripts associated with TLR signaling (Figure 1b). Other pathways include genes involved in TNF, interferon, Fc epsilon receptor, and NLR signaling pathways. Cyclin D associated events in G1, and transcriptional regulation of TP53 were also differentially bound by PU.1 binding in SP-R210_L_(DN) cells (Supplemental Figure 4).

### 4.3 Differential distribution of histone methylation marks in WT and SP-R210_L_(DN) cells

Previous studies reported that PU.1 binding is influenced by histone 3 (H3) methylation (Cheng, et al., 2013, Burda, et al., 2016, Tagore, et al., 2015). Methylation marks on lysine residues of histone 3 (H3) are associated with certain chromatin conformations; H3K4me3 with open chromatin and transcriptional activity, and H3K9me3 and H3K27me3 are typically associated with closed chromatin and thus transcriptional suppression. Profiling of by ChIP-seq showed increased H3K4 trimethylation (H3K4me3) in SP-R210_L_(DN) cells compared to WT cells (Figures 3a, 3b). Conversely, methylation marks associated with transcriptional repression, H3K9 trimethylation (H3K9me3), and H3K27 trimethylation (H3K27me3), were decreased in SP-R210_L_(DN) cells compared to WT cells (Figures 3c, 3d, 3e, and 3f). The distribution of these methylation marks was also altered around specific genomic features in SP-R210_L_(DN) cells with decreased distribution of H3K4me3 marks in promoter regions, whereas H3K4me3 was higher in introns and intergenic regions (Figure 3b). The H3K4me3 methylation pattern overlapped with 23.5% and 34.9% of PU.1 binding sites representing 1911 regions in WT cells and 10413 in SP-R210_L_(DN) cells (Figure 3g, 3h), respectively.

**Figure 3.**
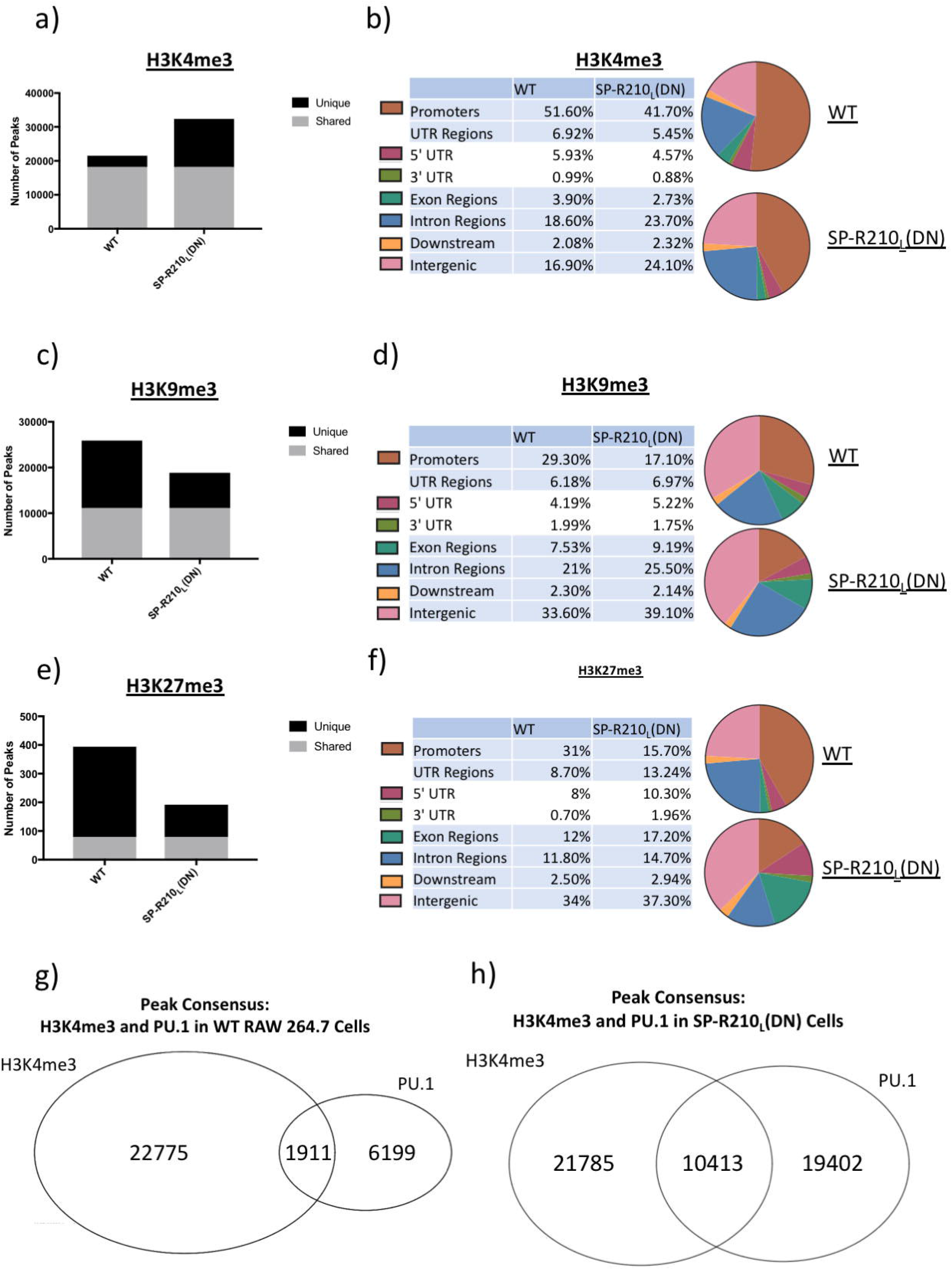
SP-R210_L_(DN) cells have altered histone methylation. **(a)** ChIP-seq of H3K4me3 was performed for both WT and SP-R210_L_(DN) cells, showing increased H3k4me3 marks in SP-R210_L_(DN) cells. **(b)** Genomic features associated with H3K4me3 marks were analyzed for both cell types, showing decreased H3K4me3 marks in promoter regions, but increased in intron and intergenic regions. This analysis was repeated for H3K9me3 **(c, d)** and H3K27me3 **(e, f)** methylation marks. For both H3K9me3 and H3K27me3, there were decreased amounts of these marks in SP-R210_L_(DN) cells, with similar changes in genomic distribution; there were decreased H3K9me3 and H3K27me3 marks in promoter regions, but increased in Exon, Intron, and Intergenic regions. Using ChIPPeakAnno, it was seen that of the H3K4me marks, only a small proportion is associated with PU.1 peaks in both WT **(g)** and SP-R210_L_(DN) **(h)** cells.

Plotting of bedgraphs on the UCSC browser was used to examine the relationship between PU.1 binding and histone methylation pattern for TLR genes, since TLR response pathways were differentially affected in SP-R210_L_(DN) cells. Concordant peaks for both PU.1 and H3K4me3 in the promoter regions of *Tlr3, 5, 6, and 9* were broad and narrow in WT and SP-R210_L_(DN) cells, respectively. However, there were additional and/or shifted peaks for PU.1 and H3K4me3 near the transcriptional start site of *Tlr2*, *Tlr5*, *Tlr6*, *Tlr9* and *Tlr13* in SP-R210_L_(DN) cells (red arrows, Supplemental Figure 5). Sharp PU.1 peaks near or further upstream the transcriptional start site for *Tlr4 and Tlr7* were not different between WT and SP-R210_L_(DN) cells, and these peaks were not concordant to H3K4me3 (Figure 4a, Supplemental Figure 5). There were no discernible PU.1 or H3K4me3 peaks for *Tlr1, Tlr3, Tlr8, Tlr11 and Tlr12* (Supplemental Figures 6). Narrow PU.1 peaks at sites distal to the promoter of *Tlr13* were discernible only in SP-R210_L_(DN) regions and these peaks were concordant to H3K4me3 (Supplemental Figure 5). Validating our immunoprecipitation experiment, PU.1 binding was identified at the known enhancer binding site upstream of the PU.1 transcriptional start site in both WT and SP-R210_L_(DN) cells (Figure 4b). A low intensity PU.1 peak in the proximal PU.1 promoter was retained in WT but not SP-R210_L_(DN) cells, whereas a broad H3K4Me3 peak was present in both (Figure 4b). By comparison, chromatin peak analysis of the *Csfr1* gene revealed broad and narrow PU.1 and H3K4Me3 peaks downstream of the first exon in WT and SP-R210_L_(DN) cells, respectively (Supplemental Figure 5). Bedgraph analysis of the *Myo18A* gene encoding SP-R210 isoforms revealed two consensus PU.1 binding motifs in the reverse complement and forward strand orientations (Figure 4c). Of these, only the intronic cis site was occupied by PU.1 in both WT and SP-R210_L_(DN) cells. An additional non-consensus PU.1 binding peak was present upstream in the same intron, suggesting indirect PU.1 binding at this site. Sequence analysis of these peaks revealed a canonical PU.1 binding sequence near the intronic peak, while a non-canonical PU.1 binding motif was identified by UCI motifmap near the upstream site (K, et al., 2011, G, et al., 2013, V, 2016). A sharp H3K4Me3 peak in the promoter region had reduced intensity in SP-R210_L_(DN) cells (Figure 4c).

**Figure 4.**
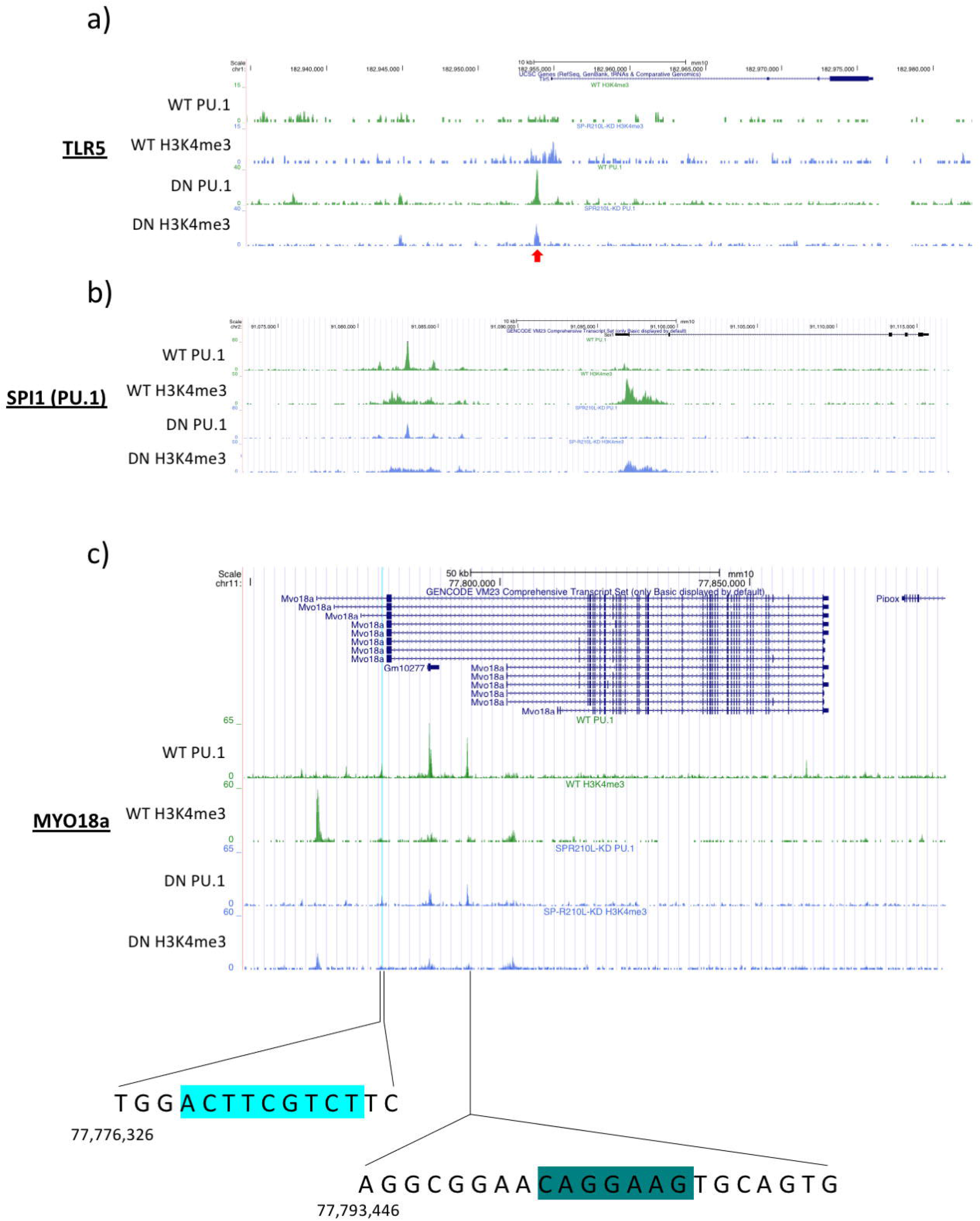
PU.1 and H3K4me3 peaks Association with Genes are Altered with SP-R210_L_(DN) cells. **(a)** Bedgraphs were generated for PU.1 and H3K4me3 ChIP and mapped to UCSC mm10 annotated genome. Viewing TLR5 on the UCSC genome browser revealed increased PU.1 and H3K4me3 binding at the promoter region of TLR5 in SP-R210_L_(DN) cells. Additional bedgraphs of TLR genes can be found in Supplemental Figure 5, Supplemental Figure 6. **(b)** PU.1 is known to bind its own enhancer region; visualizing PU.1 on the genome browser revealed PU.1 binding sites in the enhancer region of PU.1 in both WT and SP-R210_L_(DN) cells. **(c)** Investigating the Myo18A gene revealed several PU.1 peaks in both WT and SP-R210_L_(DN) cells of varying intensities. H3K4me3 peaks were found at the promoter region of Myo18a, as well as an internal start site. Highlighted in light blue is a predicted PU.1 binding site (UCI Motifmap), with the sequence depicted below. The sequence of a PU.1 peak present internal to Myo18A is also depicted. Within the sequence, a canonical PU.1 binding motif is highlighted in dark green.

### 4.4 IAV infection results in repression and redistribution of PU.1 binding and differential outcomes of cellular death and signaling responses in WT and SP-R210_L_(DN) cells

PU.1 is critical for the terminal differentiation of alveolar macrophages downstream from GM-CSF receptor signaling in the local microenvironment (Carey, et al., 2007). In turn, alveolar macrophages are essential for host survival from severe influenza A virus (IAV) infection (Halstead, et al., 2018, Sever-Chroneos, et al., 2011, Umstead, et al., 2020, Huang, et al., 2011). Raw264.7 cells have been extensively utilized as a surrogate *in vitro* model to study IAV infection in macrophages with all known IAV strains (Marvin, et al., 2017, Cline, et al., 2017). Infection of IAV in macrophages is largely abortive, whereby replication, transcription, and translation of viral genes takes place with minimal packaging and release of viral progeny (Marvin, et al., 2017, Cline, et al., 2017). Therefore, we asked whether IAV infection on alters PU.1 chromatin occupancy in WT and SP-R210_L_(DN) cells. IAV infection and lack of the L isoform did not alter PU.1 expression WT and SP-R210_L_(DN) cells (Supplemental Figure 3). ChIP-seq immunoprecipitation experiments revealed that IAV infection reduced PU.1 binding by almost 40 and 65%, respectively, compared to uninfected cells (Figure 5a, 5b). The SP-R210_L_(DN) genome had 3153 fewer PU.1 peaks compared to uninfected cells, whereas the number of PU.1 peaks in SP-R210_L_(DN) cells decreased by 7560 compared to uninfected cells (Figure 5a). Of these, 733 and 197 PU.1 peaks were uniquely associated with infection in WT and SP-R210_L_(DN) cells, respectively (Figure 5b). Mapping of these unique peak regions showed increased distribution of PU.1 peaks in proximal promoter regions by 7 and 3.3%, or a 2-fold difference between infected WT and SP-R210_L_(DN) cells, respectively. The 2-fold increase reflected lower PU.1 distribution in intronic and distal intergenic regions in WT cells compared to lower distribution in intronic regions in SP-R210_L_(DN) cells (Figure 5c). PU.1 peaks in shared genomic regions mapped to mostly macrophage function and activation pathways (e.g. Fc gamma receptor dependent phagocytosis, regulation of actin dynamics for phagocytosis, clathrin-mediated endocytosis, and signaling by RHO GTPases) in both WT and SP-R210_L_(DN) cells (Figure 5d, Supplemental Figure 8) and these were selectively suppressed after infection of SP-R210_L_(DN) cells. IAV infection, however, resulted in differential increase in association of PU.1 with genes regulating apoptotic and hyaluronan metabolism pathways, whereas binding to genes associated with proinflammatory TLR pathways decreased in WT and SP-R210_L_(DN) cells, respectively (Figure 5d, Supplemental Figure 10). Accordingly, flow cytometric experiments confirmed the pro-apoptotic phenotype of WT cells and resistance to apoptosis in response to IAV infection in WT and WT and SP-R210_L_(DN) cells, respectively (Supplemental Figure 9a) and activation of apoptotic, cell death, and sirtuin pathways (Supplemental Figure 9b). On the other hand, phosphorylation of IRF3 and IRF7 were both enhanced by 12 and 24 hrs after IAV infection in SP-R210_L_(DN) cells compared to WT (Figure 6a, b), consistent with enhancement of interferon and TLR signaling pathways. IAV infection increased Furthermore, expression of TLR7 but not TLR4 (Supplemental Figure 10) in SP-R210_L_(DN) cells compared to WT. Analysis of NFκB p65 phosphorylation revealed basal increase in the levels of serine 276 which remained elevated after infection in SP-R210_L_(DN) cells (Figure 6c), whereas phosphorylation of serine 536 was differentially induced in WT cells. Given that serine 536 phosphorylation is involved in negative regulation of NFκB signaling (Pradere, et al., 2016), we assessed phosphorylation of the upstream p65 kinase p38 (Song, et al., 2006, Schmeck, et al., 2004). Accordingly, Figure 8d shows that IAV infection induced phosphorylation of p38 in WT but not SP-R210_L_(DN) cells (Figure 6d). This difference aligns with the decrease in PU.1 binding to genes that regulate MAPK signaling (Figure 5d).

**Figure 5.**
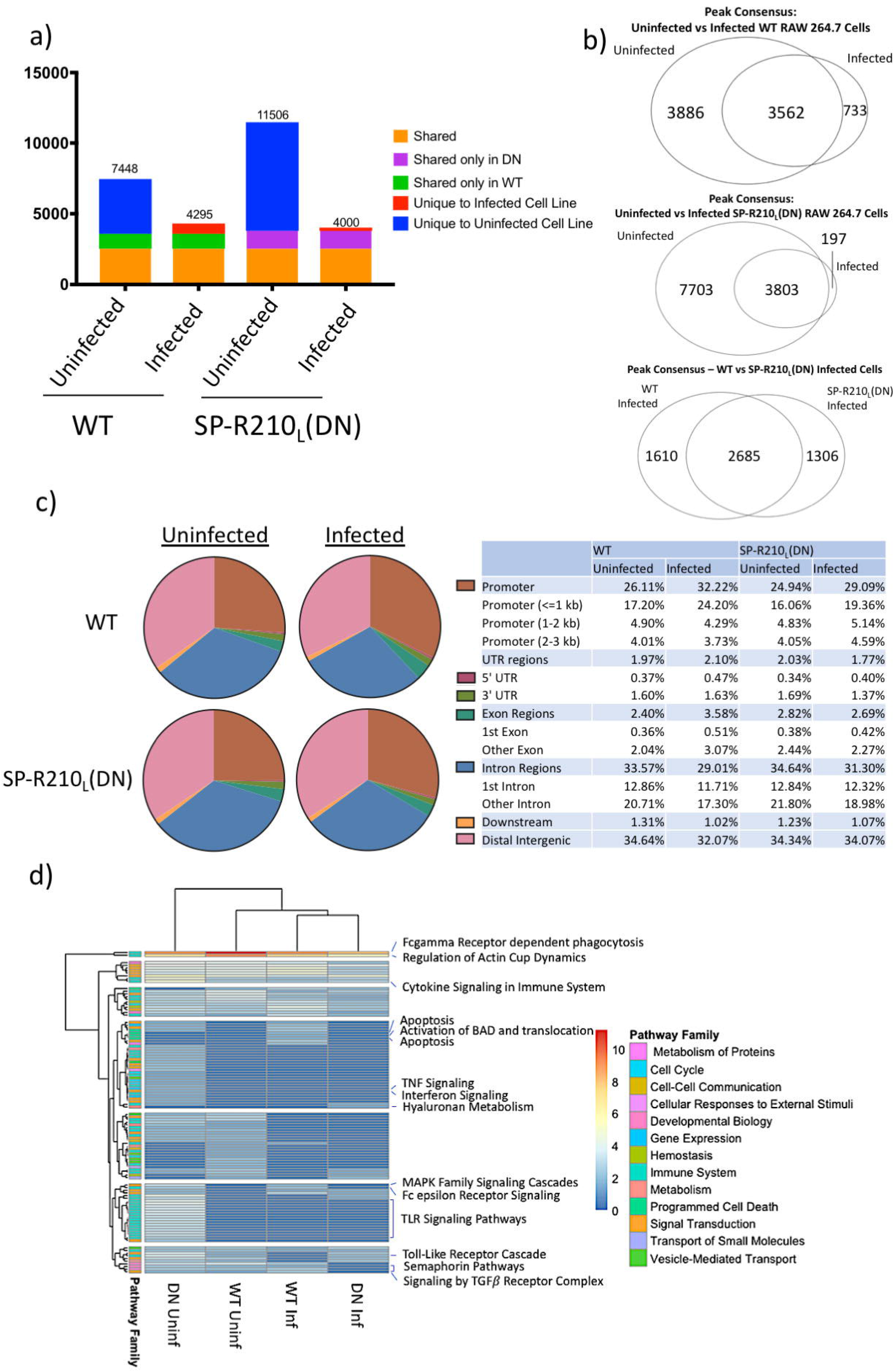
IAV infection affects PU.1 binding differently in SP-R210_L_(DN) cells than WT cells. **(a)** PU.1 peaks were compared between uninfected and infected WT and SP-R210_L_(DN) cells to determine which peaks were similar or unique to each condition; infection reduces PU.1 binding in both WT and SP-R210_L_(DN) cells. **(b)** IAV infection affects WT and SP-R210_L_(DN) cells differently; while PU.1 binding with IAV infection has many shared regions with uninfected cells, some unique PU.1 bound regions in WT and SP-R210_L_(DN) cells were identified with infection. IAV infected WT and SP-R210_L_(DN) cells showed a majority or PU.1 bound regions to be similar, but each cell type also had numerous peaks that were unique with IAV infection. **(c)** PU.1 binding was mapped to genomic features for infected and uninfected WT and SP-R210_L_(DN) cells. Mapping revealed increased distribution of PU.1 binding to promoter regions in WT and SP-R210_L_(DN) cells, with concomitant decreases in Intron and Intergenic regions. **(d)** PU.1 associated regions were mapped to Reactome pathways; Pathway enrichment scores and p-values for WT and SP-R210_L_(DN) cells were plotted in a heat map; each pathway was associated to larger Reactome pathway families.

**Figure 6.**
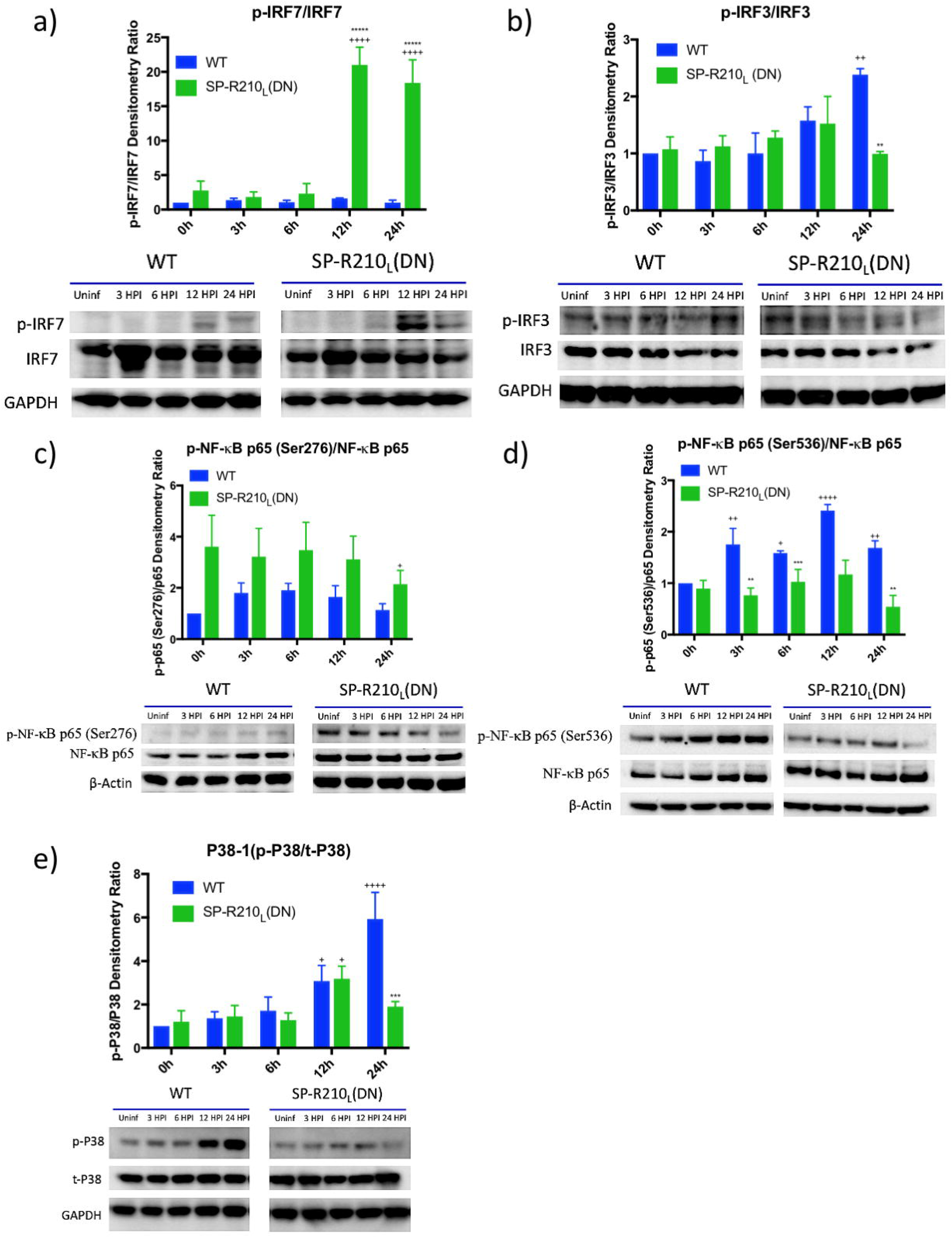
SP-R210_L_ knockdown alters phosphorylation of immune signaling molecules. **(a)** Clarified cell lysates from WT and SP-R210_L_(DN) cells infected with PR8 for 3, 6, 12, and 24 hours were probed for phosphorylated and total IRF3 (a), IRF7 (b), NFκB p65 subunit (c, d) and P38 (e). **(a)** SP-R210_L_(DN) cells show a trend towards increased baseline IRF3 phosphorylation, but less IRF3 phosphorylation at 24 HPI (n=2). **(b)** SP-R210_L_(DN) cells showed increased IRF7 phosphorylation at baseline, with significant increases at 12 and 24 HPI (n=3) **(c, d)** SP-R210_L_(DN) cells showed increased Ser276 phosphorylation **(c)**, but decreased Ser536 **(d)**, of NFκB p65 compared with WT at baseline and throughout infection. (n=3) **(e)** WT cells exhibited increased phosphorylated P38 at 24 hours post infection compared to SP-R210_L_(DN) cells (n=3). Statistical significance determined by 2-way ANOVA. **, p-value <0.005 comparing between WT and SP-R210_L_(DN); ***, p-value<0.005 comparing between WT and SP-R210_L_(DN); +, p-value <0.05 comparing between uninfected and infected time point; ++, p-value <0.005 comparing between uninfected and infected time point; +++, p-value <0.0005 comparing between uninfected and infected time point; ++++, p-value <0.00005 comparing between uninfected and infected time point

## 5 Discussion

This study explored how SP-R210 isoforms coordinate macrophage transcriptional and epigenetic regulation of macrophage function. We report that depletion of the L isoform switches macrophages to a primed state marked by heterochromatin reduction accelerating responses to inflammatory and infectious stimuli. Previous studies showed that selective disruption of SP-R210_L_ in macrophages alters activation state coupled to changes in trafficking and expression of phenotypic markers of macrophage activation and differentiation, encompassing pattern recognition, scavenger, and adhesion receptors enhancing phagocytic function and responsiveness to inflammatory and infectious stimuli extrinsic ligand-independent functions of SP-R210 that depend on relative abundance of SP-R210 isoforms (Yang, et al., 2015, Yang, et al., 2005, Sever-Chroneos, et al., 2011). Here, we extend upon these findings, showing that selective disruption of the L isoform results in broad priming and activation of antiviral response pathways and chromatin accessibility and remodeling as demonstrated by transcriptome analysis and chromatin immunoprecipitation experiments. Additionally, disruption of both isoforms appears to result in an intermediate phenotype marked by normalization of most cell-surface markers examined and moderate anti-viral response to influenza infection compared to WT and SP-R210_L_(DN) cells, supporting that the L and S isoforms modulate antagonistic extremes of macrophage. inflammatory and anti-inflammatory activation states in part through a signaling mechanism that modulates PU.1-dependent chromatin remodeling.

Transcriptome data support the novel hypothesis that L-deficient macrophages undergo metabolic, transcriptional, and epigenetic adaptation related to lipid and fatty acid uptake, utilization, and metabolism to be addressed in future studies. Thus, it is noteworthy that expression of fatty acid uptake and metabolism genes *Fads2* and *Igf1* mRNA, and cell-surface CD36 and SR-A proteins (Oishi, et al., 2017, Koundouros and Poulogiannis, 2020, Cucchi, et al., 2019, Spadaro, et al., 2017) were highly induced in uninfected SP-R210_L_(DN) cells. In contrast, deletion of both isoforms diminished cell-surface expression of CD36. Phenotypic expression level of CD36 distinguished anti-inflammatory from inflammatory macrophages in human adipose tissue based on diminished and high CD36 expression, respectively (Kralova Lesna, et al., 2016). It is also noteworthy that histone deacetylase 9 (*Hdac9*), a deacetylase for histones, transcription factors, and other signaling molecules, was highly induced in SP-R210_L_(DN) cells. HDAC9 is one of several deacetylases that promote inflammatory M1 macrophage polarization by repressing nuclear receptors and cholesterol efflux (Cao, et al., 2014). Furthermore, deacetylation of the interferon regulating kinase TBK1 by HDAC9 was shown to enhance production of Type I interferon augmenting antiviral activity (Li, et al., 2016), consistent with the basal anti-viral phenotype of the L-deficient macrophages. Accordingly, TBK1 and related antiviral control genes were induced in SP-R210_L_(DN) cells after IAV infection, whereas the infection resulted in apoptotic and sirtuin pathway involvement in WT cells. Sirtuins respond to cellular NAD+/NADH redox balance bioavailability, thereby targeting a broad range of protein substrates that drive apoptosis, DNA repair, metabolism, and inflammation in response to different cellular conditions (Zhang and Sauve, 2018).

The global transcriptome adaptation of the L-deficient cells was accompanied by differences in accessibility of the transcription factor PU.1 and chromatin remodeling as demonstrated by increased genome-wide deposition of PU.1 and epigenetic histone modification. The pattern of histone methylation marks is consistent with increased activation state of SP-R210_L_(DN) cells as indicated by increased number of genes associated with H3K4me3 compared to decreases in H3K9me3 and H3K27me3 associated chromatin, although in all cases there was marked redistribution of these trimethylated histones from promoter to non-promoter regions. As a pioneer transcription factor, PU.1 interacts with both promoter and non-promoter regions throughout the genome in both active and repressive chromatin, regulating chromatin accessibility and gene expression. PU.1 maintains chromatin at an open conformation to allow binding of stimulus-dependent transcription factors and elicit expression of macrophage activation genes by displacing nucleosomes, whereas tight control of PU.1 levels plays a critical role in the fate of hematopoietic stem cells towards myeloid or lymphocytic lineages. On the other hand, chromatin structure and composition of methylated and acetylated histones may limit PU.1 access in closed transcriptionally inactive chromatin after lineage commitment (Glass and Natoli, 2016, Hoogenkamp, et al., 2007, Imperato, et al., 2015, Petrovick, et al., 1998, Schmidt, et al., 2016, Karpurapu, et al., 2011, Liu and Ma, 2006, Rosa, et al., 2007, Celada, et al., 1996, Ha, et al., 2019, Qian, et al., 2015, Rothenberg, et al., 2019, Ghisletti, et al., 2010, van Riel and Rosenbauer, 2014, Leddin, et al., 2011, Rojo, et al., 2017, Schroder, et al., 2007, Lichanska, et al., 1999, Ross, et al., 1998, Reddy, et al., 1994). The number of PU.1 peaks in active chromatin enriched in H3K4Me3 increased five-fold in L-deficient cells, indicating marked reconfiguration of chromatin of the SP-R210_L_(DN) cell genome from a closed to an active state, although additional DNA accessibility studies are needed to validate this finding. Lack of L did not have a major impact on the overall fractional distribution of PU.1 between promoter and non-promoter binding sites in uninfected cells. Selective analysis of PU.1 bound genes revealed sharp PU.1 peaks near transcriptional start sites in SP-R210_L_(DN) cells compared to the broad heterogeneous PU.1 peaks, or a complete absence of peaks, in WT cells at proximal intronic regions downstream from the transcriptional start site in several but not all TLR genes (*Tlr3, 5, 6, 9, and 13*). Analysis of the TATA-less *Csfr1*, a known PU.1 regulated gene that encodes the M-CSF receptor, displayed similar repositioning as seen with narrow PU.1 binding peaks in the known PU.1 intronic binding sites in this gene. In contrast, the shape of the PU.1 peaks in the enhancer regions −15 to −8 kb upstream the PU.1 promoter (van Riel and Rosenbauer, 2014, Leddin, et al., 2011) were not affected. PU.1 binding consensus motifs were identified inside the first intron and upstream near the transcriptional start site on the opposite strand of the Myo18A gene, although only the intronic site was occupied in a sharp PU.1 peak and this was similar in both WT and SP-R210_L_(DN) cells. Whether this PU.1 site contributes to expression for the L isoform in mature macrophages but not macrophage precursors remains to be determined. Therefore, downregulation of the L isoform alters local positioning and complexity of intragenic PU.1 binding and association with promoter and intronic elements.

In response to IAV infection, however, there was marked expulsion of PU.1 binding, depleting PU.1 from genes affecting diverse regulatory processes in WT cells but predominantly macrophage activation genes regulating toll-like receptor signaling in SP-R210_L_(DN) cells. Assessment of bound PU.1 showed increased distribution of PU.1 promoter sites associated with activation of cellular death pathways and metabolism in WT and SP-R210_L_(DN) cells. The SP-R210_L_(DN) cells, however, retained the ability to elaborate anti-viral responses to IAV infection. These findings support the model that depletion of SP-R210_L_ results in PU.1-dependent heterochromatin reduction and basal activation of SP-R210_L-S+_ macrophages priming macrophage activation through differential regulation of p38 and NFκB pathways and activation of IRF3/7 signaling in response to influenza infection.

## 6 Conclusion

This study explored how SP-R210 isoforms coordinate macrophage transcriptional and epigenetic regulation of macrophage function. We report that depletion of the L isoform switches macrophages to a primed state marked by associated heterochromatin reduction that was accompanied by redistribution and expansion of PU.1 chromatin occupancy. This chromatin remodeling may accelerate responses to inflammatory and infectious stimuli. To this end, the mechanisms that elicit PU.1 redistribution and heterochromatin reduction in macrophages and other immune cells are only partly understood (Tagore, et al., 2015, Minderjahn, et al., 2020, McAndrew, et al., 2016). On the other hand, decline in expression of the L isoform alters SP-A binding from linear non-cooperative to binding with positive cooperative behavior (Supplemental Figure 11a and b) in WT and SP-R210_L_(DN) cells, respectively. In this regard, future studies are needed to elucidate the cross-talk between SP-A and GM-CSF (Shibata, et al., 2001, Chroneos and Shepherd, 1995, Blau, et al., 1994, Yoshida, et al., 2001), its role modulating alveolar macrophage function and local homeostasis, and whether PU.1 is the downstream effector of this interaction *in vivo*. Furthermore, it is noteworthy that a shorter 110 kDa isoform of SP-R210 was shown to interact physically with the M-CSF receptor signaling complex in human U937 monocytic cells (Cross, et al., 2004), which do not express the L isoform (Yang, et al., 2005). Therefore, future studies are needed to explore the impact of L deletion and SP-R210 deletion mutagenesis in the PU.1/M-CSF receptor signaling pathway. This may contribute to dynamic modulation of immune activation threshold of alveolar and inflammatory macrophages with different expression of SP-R210 isoforms to modulate inflammation and host resistance to infection. Our study provides the framework for further studies to delineate intrinsic and ligand-dependent mechanisms by which SP-R210 isoforms in macrophages regulate homeostatic and inflammatory functions of resident macrophage populations.

## 8 Author Contributions

EY acquired, analyzed, graphed genomic data, and wrote the manuscript. YC and EY developed and performed influenza infection assays and processed cells for RNAseq analysis. CS provided expertise for chromatin immunoprecipitation experiments. JW produced and isolated SP-R210(KO) data using CRISPR. YIA oversaw the generation of RNAseq and ChIP-derived libraries, acquisition RNAseq and ChiPseq data, provided expertise for bioinformatics analysis, and read manuscript critically. ZT and YT provided expertise and reagents for CRISPR knockout of SP-R210. TMU purified IAV, performed experiments, and edited manuscript. SD provided reagents and expertise in analysis of epigenetic data. ZC conceptualized, designed, contributed to bioinformatics analysis, led the study, and co-wrote the manuscript.

## 10 Acknowledgments

We would like to thank Nate Schaffer and Joseph Bednarzyk from the Pennsylvania State University College of Medicine Flow Cytometry Core Facility as well as the Institute of Personalized Medicine for assistance with flow cytometry and genomic processes.

## 13 Data Availability Statement

The datasets for this study can be found on GEO [].

