## Supplemental Figures and Tables for "Genomic and epigenomic adaptation in SP-R210 (Myo18A) isoform-deficient macrophages"

Supplemental Figure 1

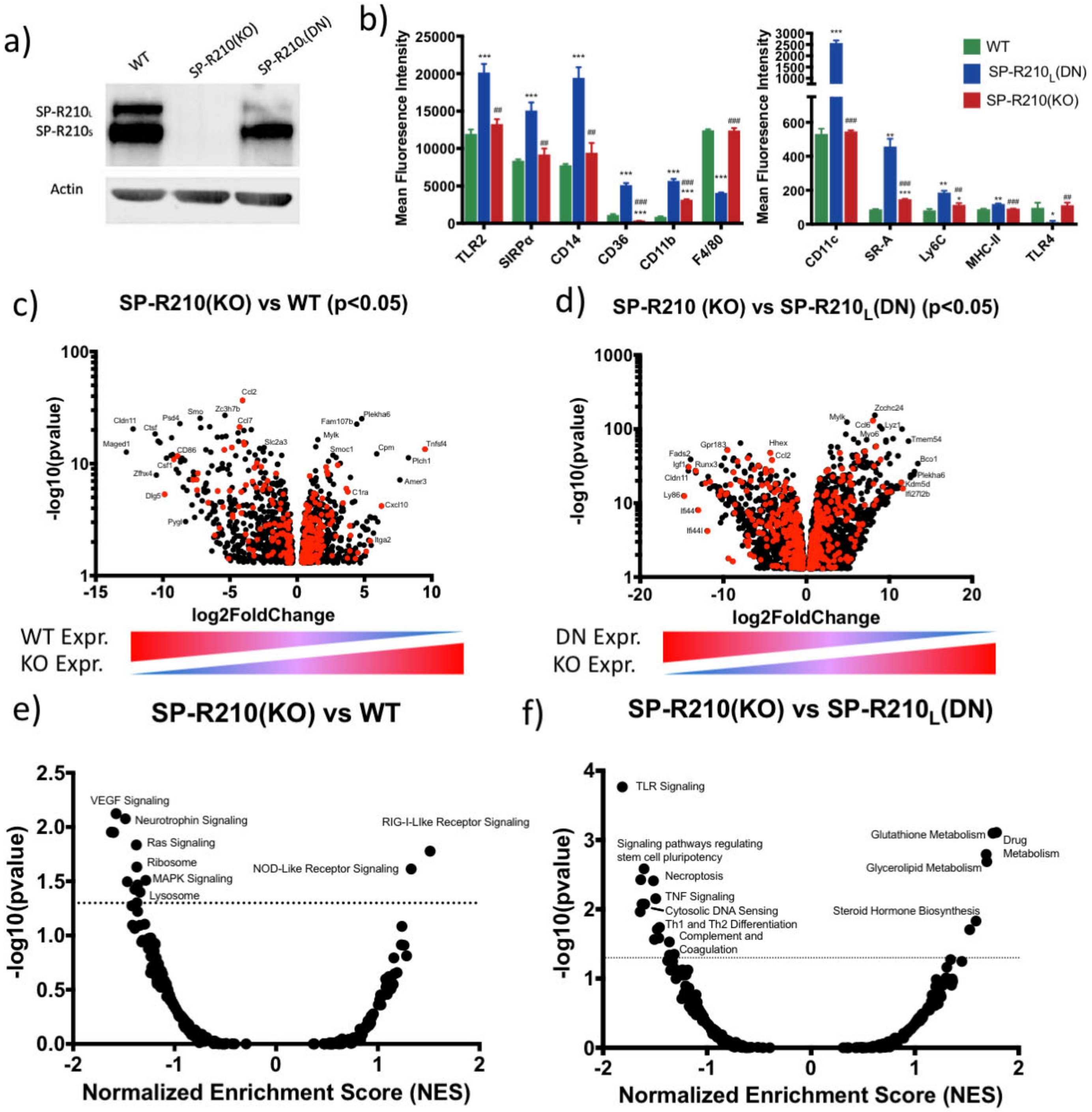


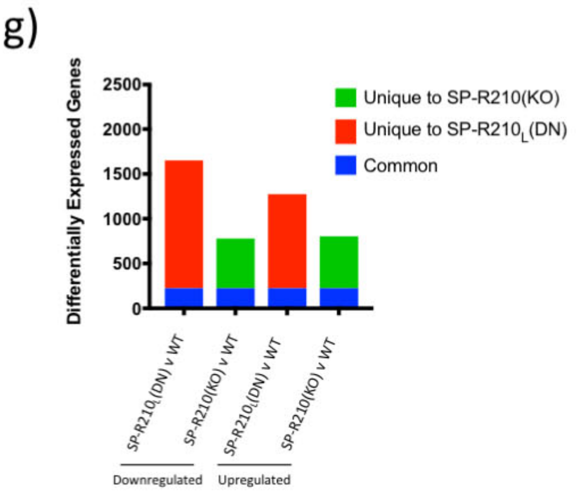


**Supplemental Figure 1 Complete Knockout of SP-R210 alters gene expression differently than knockdown of one isoform (a)** Representative Western blot of uninfected WT, SP-R210$_L$(DN) and SP-R210(KO) cells showing depletion of specific SP-R210 isoforms **(b)** 2E5 cells/well cultured overnight were stained with fluorescent cell surface marker antibodies. Data plotted is mean fluorescence intensity ± S.E. (n=3), **, adjusted p-value <0.005 compared to WT; ***, adjusted p-value <0.0005 compared to WT. **(c, d)** RNA from WT, SP-R210$_L$(DN) and SP-R210(KO) were sequenced and aligned, with differential expression of 3 replicates determined by deseq2. Differently expressed genes were filtered by p-value<0.05. Immune genes were identified by MGI Immune Genes database and labeled in red**. (e, f)** Differentially expressed genes were mapped to KEGG pathways and compared between SP-R210(KO) and WT cells **(e)** and SP-R210(KO) and SP-R210L(DN) **(f)**. **(g)** Number of differently regulated genes varied with SP-R210 isoform expression. 224 genes in common were downregulated between SP-R210$_L$(DN) and SP-R210(KO) when compared to WT. 1428 genes were uniquely downregulated in SP-R210$_L$(DN) cells and 555 genes in SP-R210(KO) when compared to WT. 1049 genes were uniquely upregulated in SP-R210$_L$(DN) and 580 genes in SP-R210(KO) when compared to WT.

Supplemental Figure 2

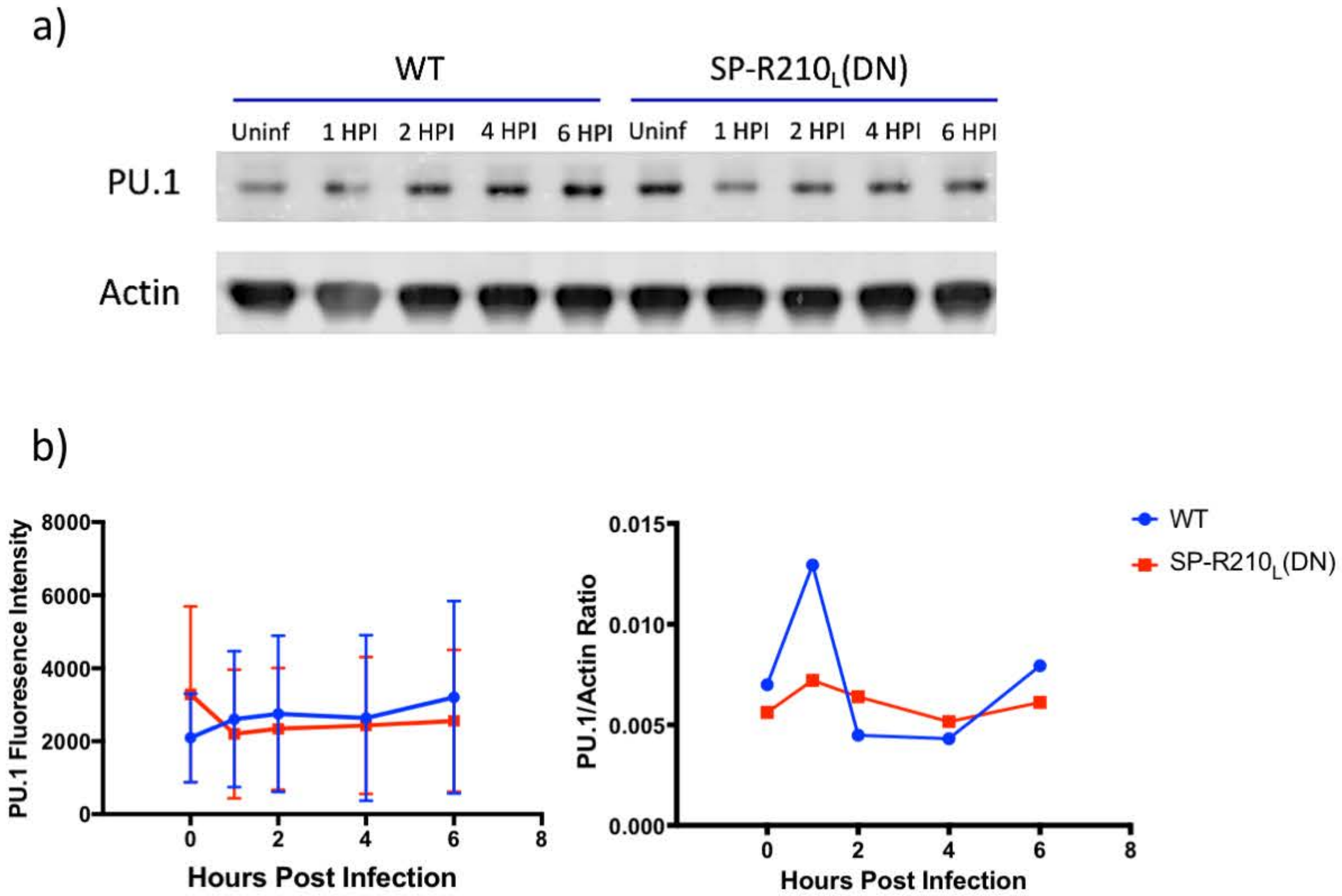


**Supplemental Figure 2. Western blot analysis of PU.1 expression before and after IAV infection.** WT and SP-R210$_L$(DN) cells were plated at 2E5 cells/well, and infected with MOI4 of IAV for indicated period of time. Cells were lysed and cell pellets were ran on 4-12% Bis-Tris gels, and transferred for western blot analysis. (a) Representative western blot image showing slight increases in PU.1 levels over time in WT and SP-R210$_L$(DN) cells over time. (b) Quantitative assessment of PU.1 fluorescence intensity of n=4 replicate experiments, as well as of ratio of PU.1 fluorescence to Actin fluorescence. Data suggests trends towards slight increases in PU.1 levels, but increases were not significant at all time points in all cell lines.

Supplemental Figure 3

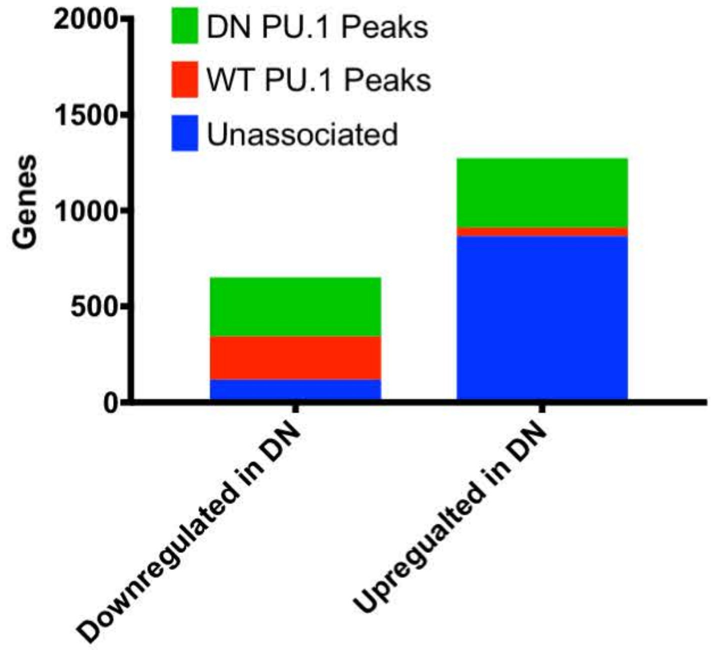


**Supplemental Figure 3. Downregulated genes in SP-R210$_L$(DN) cells are more highly associated with changes in PU.1 binding.** Upregulated and downregulated genes from DESeq2 results of SP-R210L(DN) vs WT cells were filtered by genes associated with PU.1 peaks unique to either WT or SP-R210$_L$(DN) cells. Of the 1273 genes upregulated in SP-R210$_L$(DN) cells, 44 were associated with unique PU.1 binding sites in WT cells, while 363 were associated with unique PU.1 binding regions in SP-R210$_L$(DN) cells. Of the 656 downregulated genes, 226 were associated with unique PU.1 binding in WT cells and 308 were associated with unique PU.1 binding in SP-R210$_L$(DN) cells.

### Supplemental Figure 4

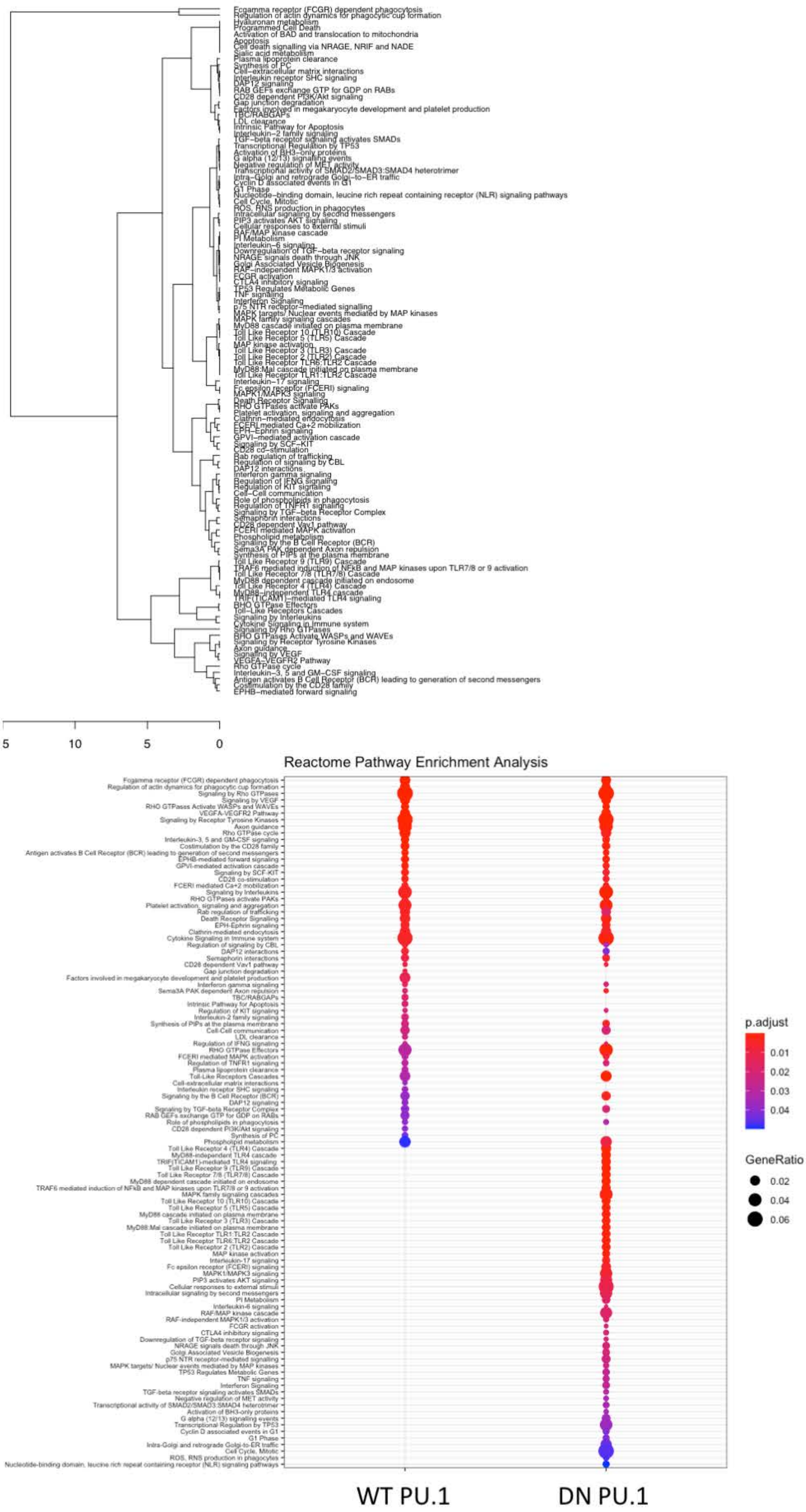


**Supplemental Figure 4. Differentially Regulated Pathways by PU.1 between SP-R210L(DN) and WT RAW cells. (A)** Dendrogram plot showing list of differentially regulated pathways by PU.1 between WT and SP-R210$_L$(DN) cells. Dendrogram plot lists pathways and is associated with the heatmap in Figure 4c. **(B)** Dotplot generated from clusterProlifer package shows Gene Ratio (proportion of genes found in analysis compared to total genes in pathway) and p-value of Reactome Pathway Analysis of PU.1 binding sites between WT and SP-R210$_L$(DN) cells.

### Supplemental Figure 5

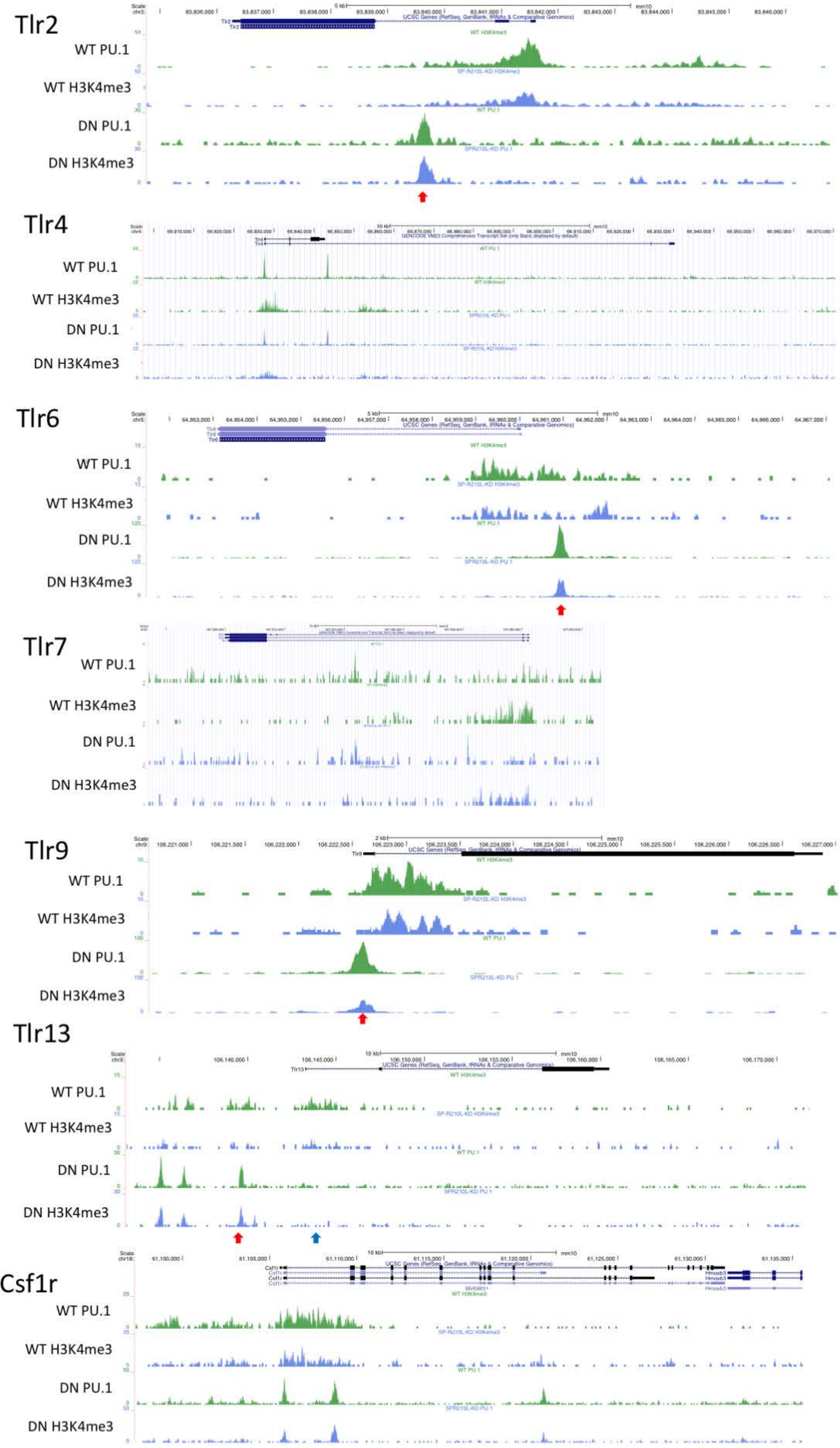


**Supplemental Figure 5. Genomic Distribution of PU.1 differs between SP-R210L(DN) and WT RAW Cells in some TLR genes.** Peaks from one representative ChIP experiment for PU.1 and H3K4me3 was plotted on the mm10 genome via UCSC genome browser. Bedgraphs for Tlr2, Tlr4, Tlr6, Tlr7, Tlr9, and Tlr13 are represented here. Red arrows indicate regions of genome with differential PU.1 and H3k4me3 peaks in SP-R210$_L$(DN) cells compared to WT cells. Csf1r also shows differences in PU.1 and H3k4me3 binding are seen in the promoter region for WT cells compared to SP-R210$_L$(DN) cells.

### Supplemental Figure 6

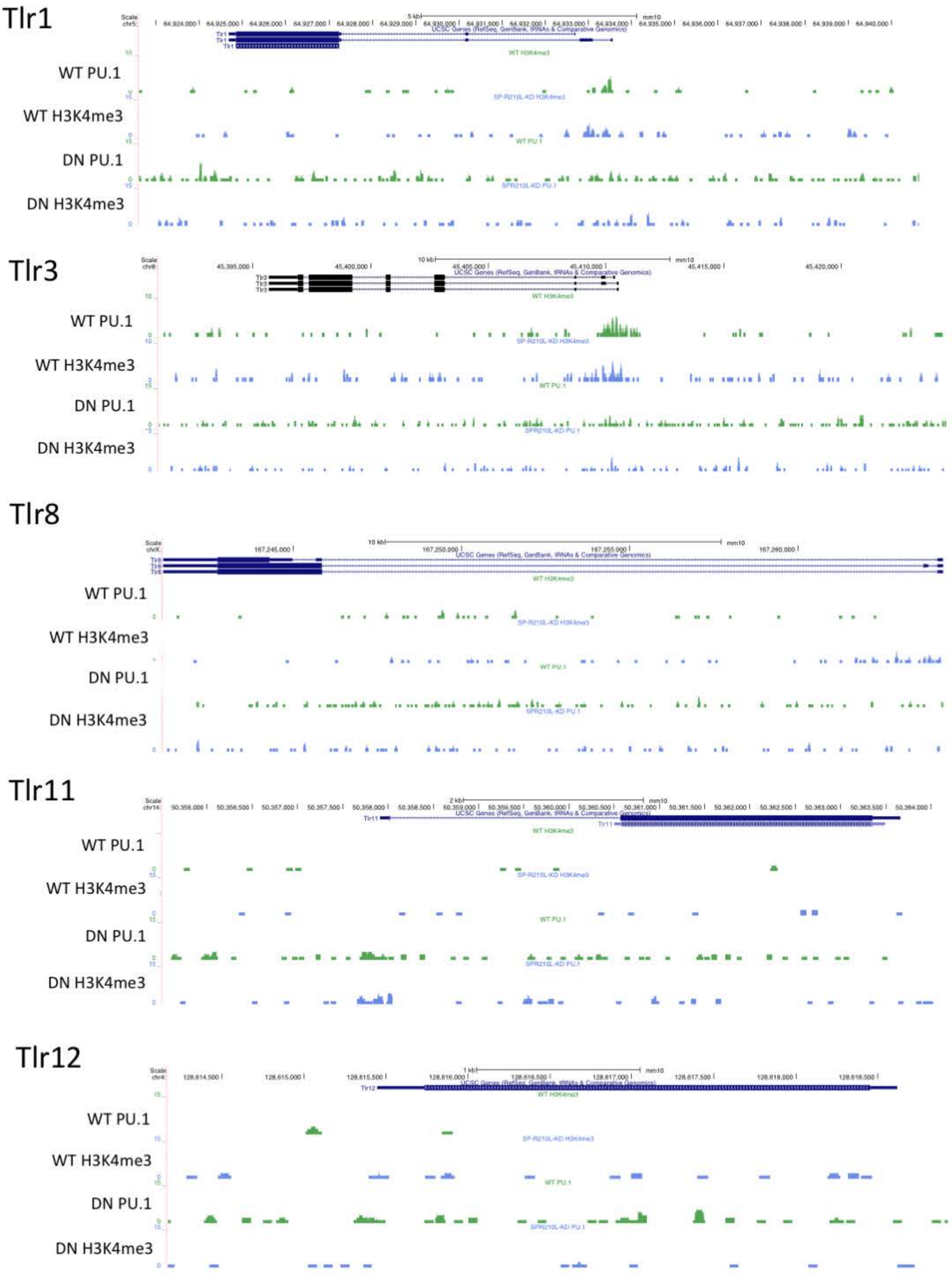


**Supplemental Figure 6. Some TLR genes do not have appreciable differences in genomic distribution of PU.1 between SP-R210L(DN) and WT RAW cells.** Peaks from one representative ChIP experiment for PU.1 and H3K4me3 was plotted on the mm10 genome via UCSC genome browser. Bedgraphs for Tlr1, Tlr3, Tlr8, Tlr11, and Tlr12 are represented here.

### Supplemental Figure 7

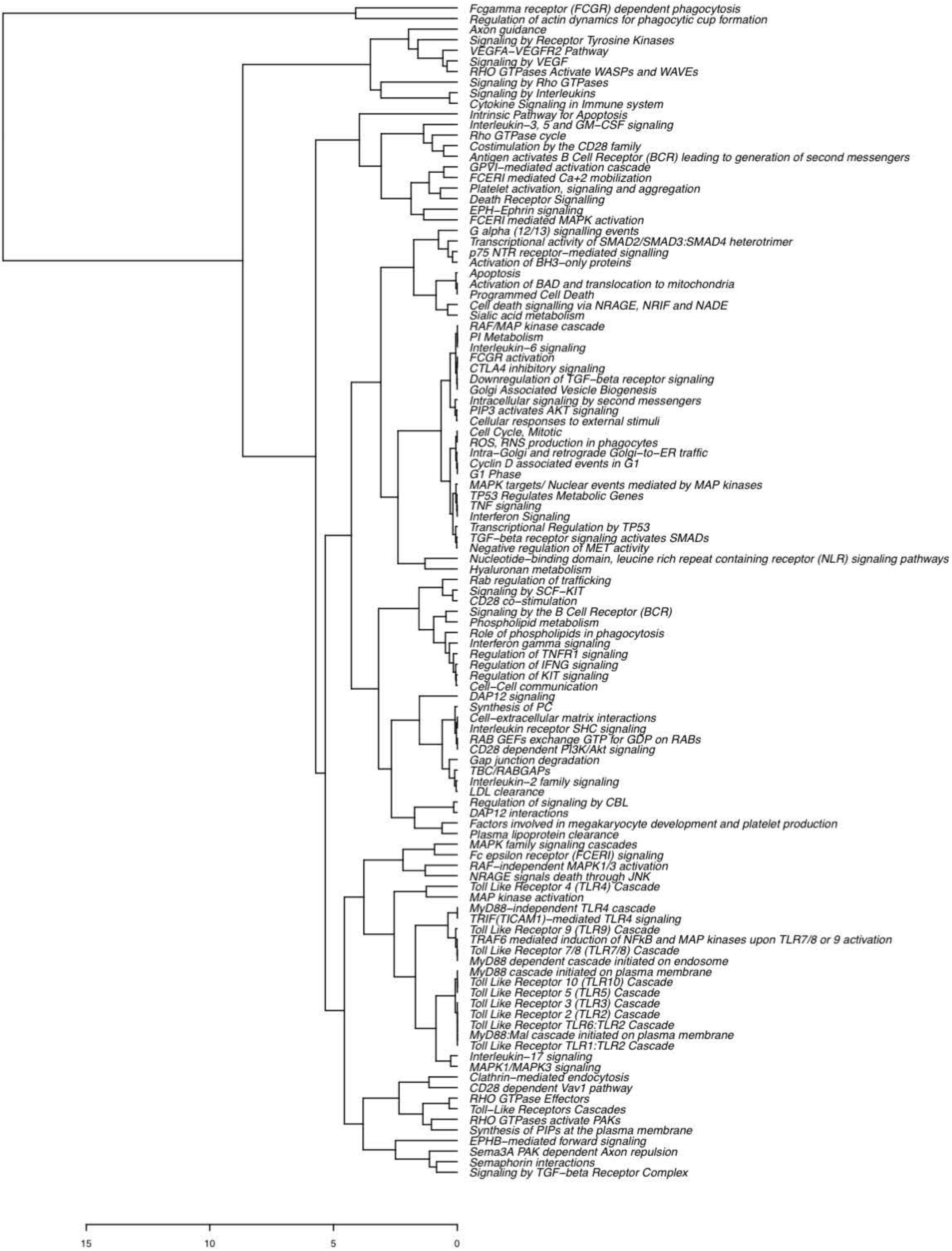


**Supplemental Figure 7. Differentially Regulated Pathways by PU.1 during Infection for WT and SP-R210$_L$(DN) Cells.** Dendrogram plot showing list of differentially regulated pathways by PU.1 between all conditions of infected and uninfected WT and SP-R210$_L$(DN) cells. Dendrogram labeling corresponds to heatmap from Figure 7d.

### Supplemental Figure 8

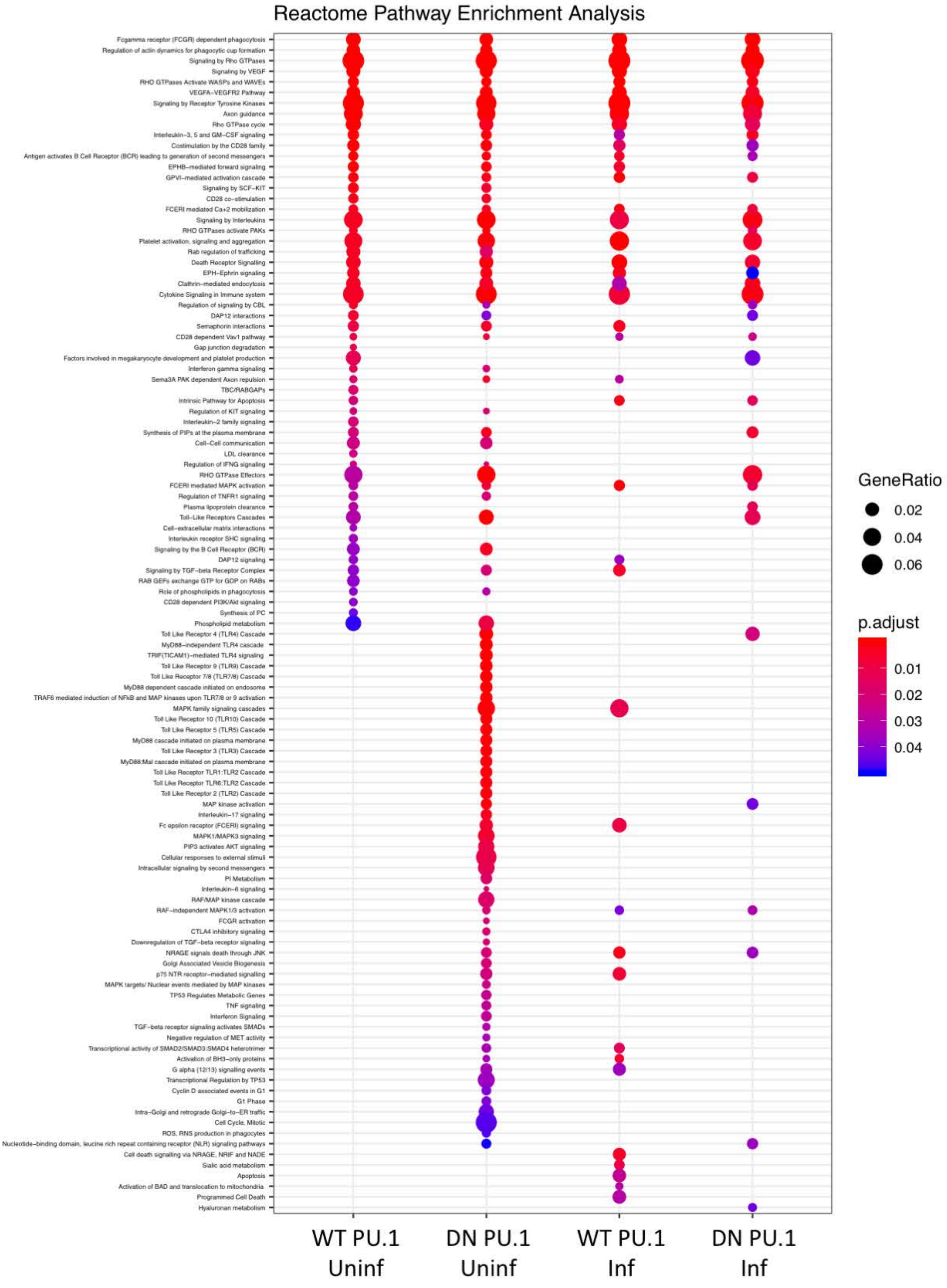


**Supplemental Figure 8. Dotplot Reveals Clustering of TNF pathways in SP-R210$_L$(DN) cells that disappears with IAV infection.** Pathways were determined and plotted from PU.1 peaks for each condition using the clusterProfiler package. Dotplot shows Gene Ratio (proportion of genes found in analysis compared to total genes in pathway) and p-value of Reactome Pathways for each condition.

Supplemental Figure 9

a)

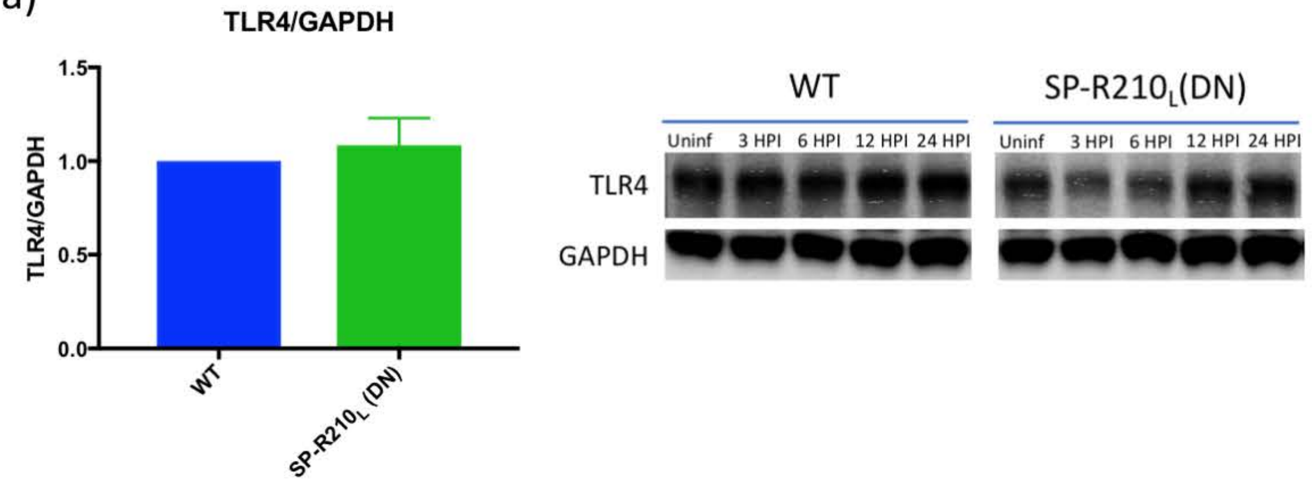


b)

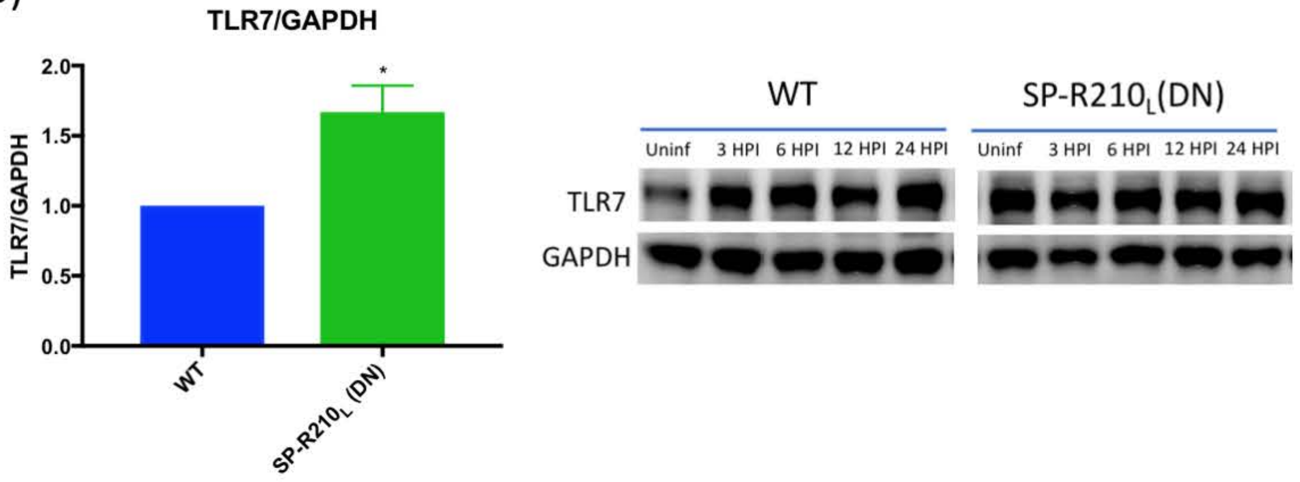


**Supplemental Figure 9. TLR4 and TLR7 expression differences between WT and SP-R210$_L$(DN) cells (a)** TLR4/GAPDH densitometry ratio of uninfected cells show no significant differences between the two cell types (n=8). Included representative western blot images of TLR4 and GAPDH with IAV infection (MOI4). **(b** TLR7/GAPDH densitometry ratio of uninfected cells shows a significant increase in TLR7 expression in SP-R210$_L$(DN) cells (n=6). Included representative western blot images of TLR4 and GAPDH with IAV infection (MOI4). *, adjusted p-value <0.05

Supplemental Figure 10

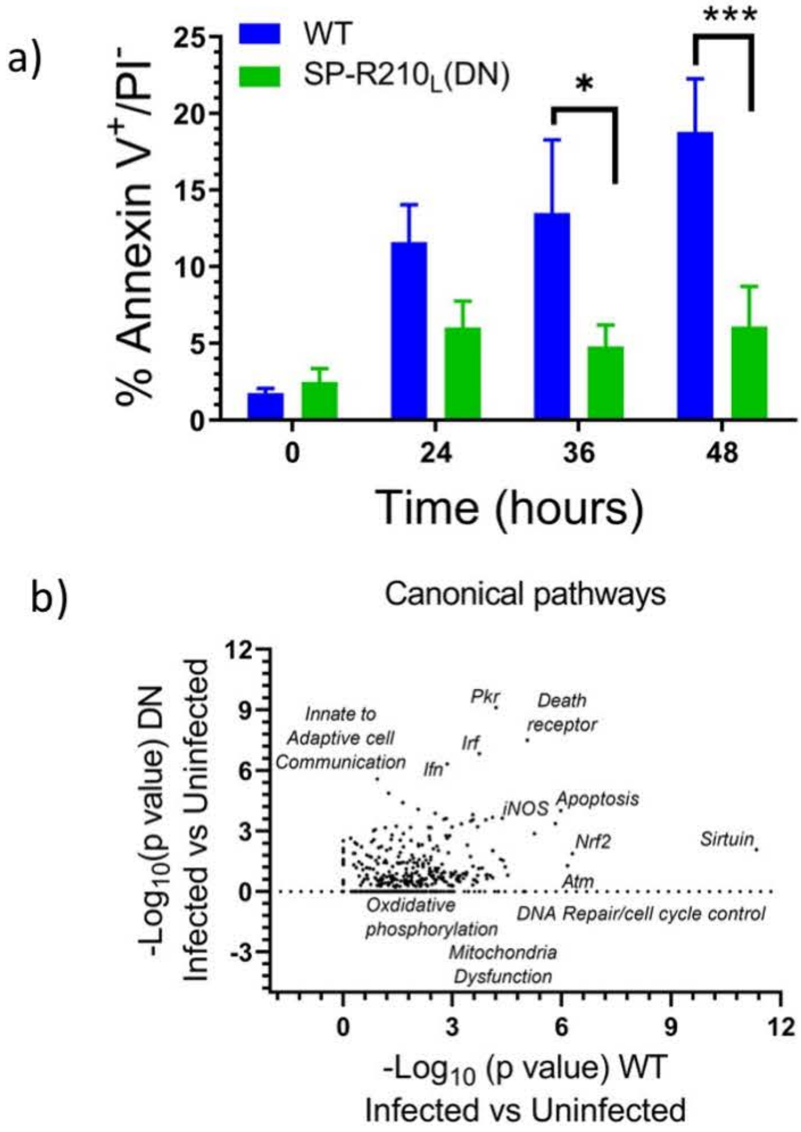


**Supplemental Figure 10. Lack of SP-R210$_L$ suppresses IAV apoptosis and activation of apoptotic, sirtuin, and DNA damage response pathways in SP-R210$_L$(DN) cells. (a)** Assessment of apoptotic Annexin V/PI- WT and SP-R210$_L$(DN) cells over time after IAV infection. **(b)** Ingenuity canonical pathway analysis of RNAseq data 24 after IAV infection.

Supplemental Figure 11

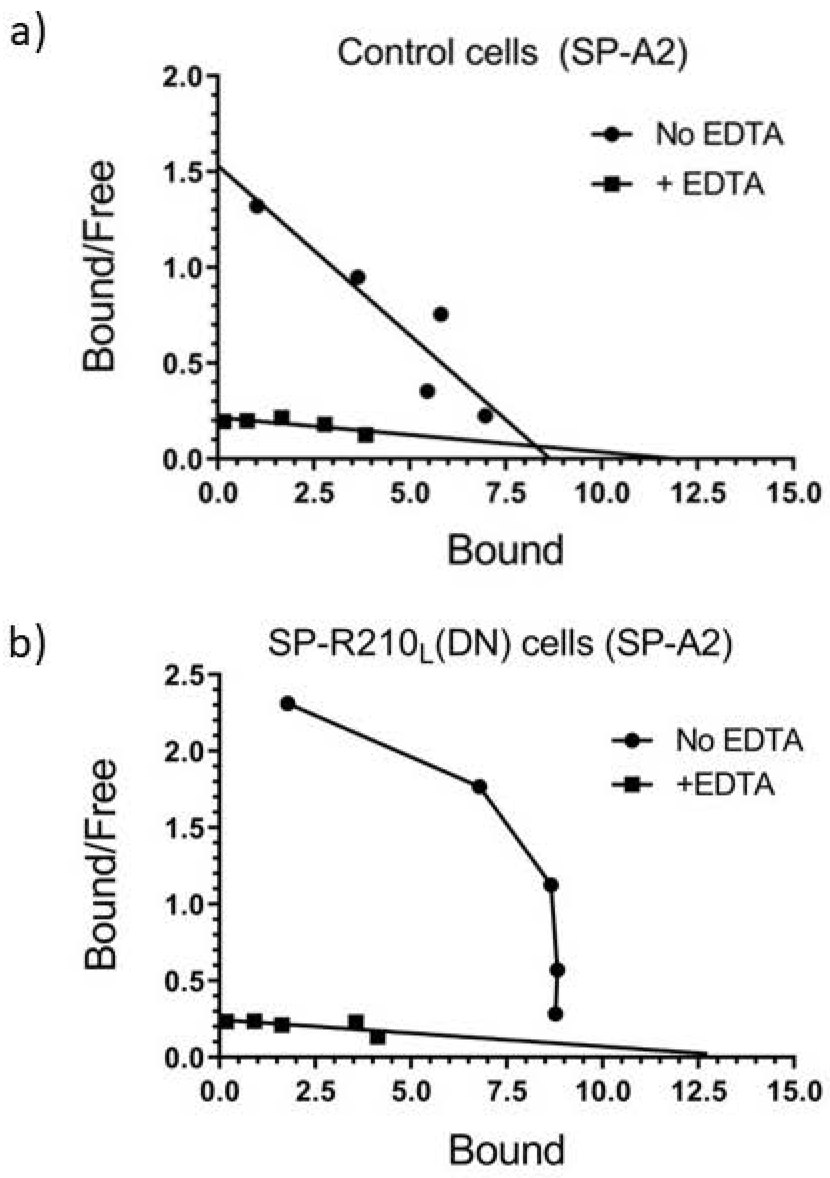


**Supplemental Figure 11. Disruption of SP-R210$_L$ alters SP-A binding characteristics. (a-b)** Scatchard analysis of human human SP-A2 binding to WT and SP-R210$_L$(DN) cells**.** Binding assays were performed as described in detail recently [1].

1. Nalian, A., et al., *Structural and Functional Determinants of Rodent and Human Surfactant Protein A: A Synthesis of Binding and Computational Data.* Front Immunol, 2019. **10**: p. 2613.

### Supplemental Table 1

## a) SP-R210$_L$(DN) vs WT

| Gene Name | baseMean | log2FoldChange | lfcSE | stat | pvalue | padj | log10(pvalue) |
|---|---|---|---|---|---|---|---|
| Top 20 Upregulated | | | | | | | |
| Ifi44l | 203.2227 | 11.4890 | 2.9747 | 3.8623 | 0.00011233 | 0.00208237 | 3.9495 |
| 2510009E07Rik | 203.7269 | 11.4880 | 1.3526 | 8.4931 | 2.01E-17 | 2.51E-15 | 16.6963 |
| Ly86 | 1421.6730 | 10.8861 | 1.7479 | 6.2280 | 4.72E-10 | 2.43E-08 | 9.3258 |
| Igf1 | 984.2160 | 10.3633 | 0.6620 | 15.6536 | 3.14E-55 | 2.68E-52 | 54.5028 |
| Cbr3 | 82.3391 | 10.1883 | 1.3011 | 7.8304 | 4.86E-15 | 4.45E-13 | 14.3130 |
| Panx1 | 76.4147 | 10.0799 | 1.2479 | 8.0777 | 6.60E-16 | 7.00E-14 | 15.1803 |
| Runx3 | 539.4463 | 10.0703 | 0.7789 | 12.9284 | 3.11E-38 | 1.69E-35 | 37.5069 |
| Gtpbp10 | 72.1738 | 9.9972 | 1.2485 | 8.0076 | 1.17E-15 | 1.19E-13 | 14.9318 |
| Ifi44 | 441.1676 | 9.9928 | 2.0913 | 4.7782 | 1.77E-06 | 4.99E-05 | 5.7523 |
| Cxcl10 | 69.6985 | 9.8265 | 1.5574 | 6.3098 | 2.79E-10 | 1.48E-08 | 9.5537 |
| Fgf13 | 58.1412 | 9.6834 | 2.0058 | 4.8276 | 1.38E-06 | 3.99E-05 | 5.8596 |
| Smagp | 56.0522 | 9.5978 | 1.2465 | 7.6996 | 1.37E-14 | 1.19E-12 | 13.8648 |
| Fam135a | 178.8361 | 9.4934 | 1.1274 | 8.4206 | 3.75E-17 | 4.45E-15 | 16.4264 |
| Oaf | 50.7069 | 9.4898 | 1.3632 | 6.9614 | 3.37E-12 | 2.21E-10 | 11.4725 |
| Kcnk5 | 50.6760 | 9.4716 | 1.3334 | 7.1035 | 1.22E-12 | 8.39E-11 | 11.9150 |
| Fads2 | 2937.5049 | 9.2749 | 0.3577 | 25.9304 | 3.02E-148 | 5.42E-144 | 147.5197 |
| N4bp3 | 43.7184 | 9.2618 | 1.3028 | 7.1090 | 1.17E-12 | 8.09E-11 | 11.9322 |
| Klhl23 | 42.9972 | 9.2431 | 1.2614 | 7.3278 | 2.34E-13 | 1.78E-11 | 12.6308 |
| Kif17 | 41.9871 | 9.1936 | 1.2650 | 7.2675 | 3.66E-13 | 2.69E-11 | 12.4363 |
| Hdac9 | 126.6371 | 9.0052 | 1.2061 | 7.4663 | 8.25E-14 | 6.54E-12 | 13.0838 |
| Top 20 Downregulated | | | | | | | |
| Ifi27l2b | 522.1791 | -10.3525 | 1.4111 | -7.3364 | 2.19E-13 | 1.67E-11 | 12.6589 |
| Gm48277 | 91.6432 | -10.3852 | 1.5601 | -6.6567 | 2.80E-11 | 1.67E-09 | 10.5528 |
| Uty | 190.0773 | -10.5056 | 1.3051 | -8.0495 | 8.31E-16 | 8.66E-14 | 15.0803 |
| Acot6 | 161.6218 | -10.5117 | 1.2363 | -8.5027 | 1.85E-17 | 2.35E-15 | 16.7324 |
| Rnf217 | 214.7451 | -10.5215 | 1.3373 | -7.8678 | 3.61E-15 | 3.37E-13 | 14.4426 |
| Prima1 | 112.3660 | -10.6061 | 1.2653 | -8.3822 | 5.19E-17 | 6.05E-15 | 16.2844 |
| Acot4 | 95.7875 | -10.6431 | 1.2855 | -8.2794 | 1.24E-16 | 1.40E-14 | 15.9072 |
| Col5a1 | 1565.7028 | -11.1174 | 0.6942 | -16.0141 | 1.02E-57 | 1.01E-54 | 56.9922 |
| Kdm5d | 300.6335 | -11.1455 | 1.2554 | -8.8778 | 6.82E-19 | 9.48E-17 | 18.1662 |
| Ddx3y | 503.0664 | -11.2852 | 1.2725 | -8.8682 | 7.43E-19 | 1.02E-16 | 18.1290 |
| Prss35 | 15299.5984 | -11.3116 | 0.5408 | -20.9154 | 3.88E-97 | 1.39E-93 | 96.4116 |
| Marcks | 250.2418 | -11.6902 | 1.3767 | -8.4918 | 2.04E-17 | 2.52E-15 | 16.6914 |
| Fxyd6 | 407.1264 | -12.1110 | 1.3390 | -9.0451 | 1.50E-19 | 2.23E-17 | 18.8251 |
| Rasd2 | 286.3148 | -12.1703 | 1.3373 | -9.1009 | 8.96E-20 | 1.40E-17 | 19.0476 |
| Eif2s3y | 707.7215 | -12.3460 | 1.2405 | -9.9528 | 2.45E-23 | 4.83E-21 | 22.6104 |
| Tmem54 | 5517.4632 | -12.4190 | 0.6979 | -17.7940 | 7.87E-71 | 1.57E-67 | 70.1043 |
| Maged1 | 352.7385 | -12.5319 | 1.7316 | -7.2370 | 4.59E-13 | 3.32E-11 | 12.3385 |
| Grb7 | 753.5820 | -12.7036 | 1.2290 | -10.3363 | 4.83E-25 | 1.06E-22 | 24.3163 |
| Ptprb | 915.0923 | -12.7929 | 1.2019 | -10.6439 | 1.86E-26 | 4.63E-24 | 25.7302 |
| Bco1 | 3972.7938 | -13.0311 | 1.0900 | -11.9550 | 6.11E-33 | 2.15E-30 | 32.2137 |

## b) SP-R210(KO) vs WT

| Gene Name | baseMean | log2FoldChange | lfcSE | stat | pvalue | padj | log10(pvalue) |
|---|---|---|---|---|---|---|---|
| Top 20 Upregulated | | | | | | | |
| Tnfsf4 | 51.2363 | 9.5011 | 1.2541 | 7.5758 | 3.57E-14 | 2.08E-11 | 13.4475 |
| Plch1 | 75.7269 | 8.2680 | 1.1975 | 6.9046 | 5.04E-12 | 1.87E-09 | 11.2980 |
| Amer3 | 14.0401 | 7.6336 | 1.4149 | 5.3950 | 6.85E-08 | 1.20E-05 | 7.1641 |
| Cxcl10 | 69.6985 | 6.2654 | 1.5678 | 3.9962 | 6.44E-05 | 0.00452121 | 4.1914 |
| 4732419C18Rik | 4.7554 | 5.9875 | 1.8103 | 3.3075 | 0.00094125 | 0.0414741 | 3.0263 |
| Cpm | 47.5577 | 5.9107 | 0.8200 | 7.2078 | 5.69E-13 | 2.62E-10 | 12.2451 |
| 9130208D14Rik | 4.4926 | 5.7802 | 1.6672 | 3.4669 | 0.00052641 | 0.02692521 | 3.2787 |
| Megf10 | 4.6126 | 5.6736 | 1.9701 | 2.8799 | 0.00397818 | 0.11895766 | 2.4003 |
| Ap1m2 | 7.9011 | 5.5251 | 1.5895 | 3.4760 | 0.00050888 | 0.0262592 | 3.2934 |
| Anks4b | 3.1885 | 5.4957 | 2.2229 | 2.4723 | 0.01342382 | 0.26929219 | 1.8721 |
| Itga2 | 3.2986 | 5.4196 | 2.0753 | 2.6115 | 0.00901423 | 0.20742839 | 2.0451 |
| Npas2 | 5.3874 | 5.4119 | 1.5396 | 3.5151 | 0.00043961 | 0.02327546 | 3.3569 |
| Urah | 2.8907 | 5.3160 | 2.0168 | 2.6359 | 0.00839228 | 0.19973621 | 2.0761 |
| Pklr | 8.9962 | 5.2484 | 1.7407 | 3.0152 | 0.00256835 | 0.08461057 | 2.5903 |
| Gm9870 | 4.1712 | 5.1790 | 1.5564 | 3.3275 | 0.00087636 | 0.03929964 | 3.0573 |
| Il12a | 2.4755 | 5.1296 | 2.2472 | 2.2827 | 0.02244826 | NA | 1.6488 |
| Gm18329 | 5.2616 | 5.0942 | 1.6980 | 3.0001 | 0.00269915 | 0.0875996 | 2.5688 |
| 1700020N01Rik | 2.3588 | 5.0605 | 2.3468 | 2.1563 | 0.03105763 | NA | 1.5078 |
| Itih4 | 2.5263 | 4.9766 | 2.2933 | 2.1701 | 0.0299997 | 0.43430496 | 1.5229 |
| Gm43059 | 4.5420 | 4.9724 | 2.0081 | 2.4762 | 0.01328036 | 0.26775752 | 1.8768 |
| Top 20 Downregulated | | | | | | | |
| Zc4h2 | 41.8326 | -8.5545 | 1.2454 | -6.8690 | 6.47E-12 | 2.31E-09 | 11.1894 |
| Man1a | 117.4970 | -8.6182 | 1.2784 | -6.7413 | 1.57E-11 | 5.18E-09 | 10.8042 |
| Tcaf1 | 57.8217 | -8.6393 | 1.2718 | -6.7927 | 1.10E-11 | 3.77E-09 | 10.9585 |
| Ppp1r9a | 144.1616 | -8.6742 | 1.3154 | -6.5943 | 4.27E-11 | 1.29E-08 | 10.3692 |
| Psd4 | 464.0704 | -8.7144 | 0.8727 | -9.9859 | 1.76E-23 | 6.15E-20 | 22.7551 |
| Gm15398 | 36.0908 | -8.8201 | 1.5118 | -5.8342 | 5.40E-09 | 1.14E-06 | 8.2673 |
| Cd86 | 190.6453 | -8.8913 | 1.2630 | -7.0398 | 1.92E-12 | 7.48E-10 | 11.7156 |
| Gprin3 | 26.6079 | -8.9880 | 1.4806 | -6.0707 | 1.27E-09 | 3.01E-07 | 8.8950 |
| Csf1 | 270.2009 | -9.1776 | 1.3549 | -6.7737 | 1.26E-11 | 4.22E-09 | 10.9013 |
| Zfp358 | 60.5509 | -9.2625 | 1.2962 | -7.1459 | 8.94E-13 | 4.01E-10 | 12.0487 |
| Vopp1 | 123.3102 | -9.3464 | 1.0887 | -8.5851 | 9.08E-18 | 8.82E-15 | 17.0420 |
| Zfhx3 | 46.5186 | -9.3903 | 1.3258 | -7.0825 | 1.42E-12 | 5.76E-10 | 11.8489 |
| Ptgs2 | 355.8014 | -9.7316 | 1.4152 | -6.8763 | 6.14E-12 | 2.24E-09 | 11.2115 |
| Dlg5 | 48.9642 | -9.8710 | 2.1559 | -4.5785 | 4.68E-06 | 0.00049648 | 5.3295 |
| Lacc1 | 62.0930 | -10.2160 | 1.2562 | -8.1322 | 4.22E-16 | 3.21E-13 | 15.3752 |
| Sdc1 | 588.3062 | -10.3561 | 1.2511 | -8.2779 | 1.25E-16 | 1.10E-13 | 15.9017 |
| Zfhx4 | 118.8838 | -10.4851 | 1.8409 | -5.6955 | 1.23E-08 | 2.42E-06 | 7.9100 |
| Ctsf | 961.5254 | -10.5731 | 1.1810 | -8.9526 | 3.47E-19 | 4.34E-16 | 18.4596 |
| Cldn11 | 1221.9780 | -12.2107 | 1.2944 | -9.4332 | 3.98E-21 | 6.96E-18 | 20.4003 |
| Maged1 | 352.7385 | -12.7204 | 1.7316 | -7.3458 | 2.04E-13 | 1.05E-10 | 12.6893 |

## c) SP-R210(KO) SP-R210$_L$(DN)

| Gene Name | baseMean | log2FoldChange | lfcSE | stat | pvalue | padj | log10(pvalue) |
|---|---|---|---|---|---|---|---|
| Top 20 Upregulated | | | | | | | |
| Bco1 | 3972.7938 | 13.4246 | 1.0900 | 12.3164 | 7.40E-35 | 2.61E-32 | 34.1310 |
| Ptprb | 915.0923 | 13.0129 | 1.2018 | 10.8279 | 2.54E-27 | 5.18E-25 | 26.5951 |
| Plekha6 | 486.6907 | 12.9457 | 1.2568 | 10.3009 | 6.98E-25 | 1.29E-22 | 24.1561 |
| Eif2s3y | 707.7215 | 12.7050 | 1.2403 | 10.2434 | 1.27E-24 | 2.28E-22 | 23.8971 |
| Grb7 | 753.5820 | 12.5464 | 1.2290 | 10.2089 | 1.81E-24 | 3.16E-22 | 23.7424 |
| Ddx3y | 503.0664 | 12.5385 | 1.2720 | 9.8570 | 6.39E-23 | 1.02E-20 | 22.1943 |
| Tmem54 | 5517.4632 | 12.3056 | 0.6979 | 17.6319 | 1.40E-69 | 2.14E-66 | 68.8534 |
| Ifi27l2b | 522.1791 | 11.6380 | 1.4107 | 8.2501 | 1.58E-16 | 1.36E-14 | 15.8005 |
| Prss35 | 15299.5984 | 11.5228 | 0.5408 | 21.3064 | 9.90E-101 | 3.03E-97 | 100.0044 |
| Kdm5d | 300.6335 | 11.4414 | 1.2551 | 9.1158 | 7.81E-20 | 9.43E-18 | 19.1075 |
| Fxyd6 | 407.1264 | 11.2366 | 1.3391 | 8.3912 | 4.81E-17 | 4.46E-15 | 16.3177 |
| Rnf217 | 214.7451 | 11.0618 | 1.3367 | 8.2754 | 1.28E-16 | 1.12E-14 | 15.8927 |
| Cyp21a1 | 1647.6815 | 11.0402 | 0.7930 | 13.9217 | 4.68E-44 | 2.77E-41 | 43.3298 |
| Elavl4 | 142.9536 | 10.8995 | 1.2696 | 8.5847 | 9.11E-18 | 9.24E-16 | 17.0406 |
| Uty | 190.0773 | 10.7619 | 1.3046 | 8.2489 | 1.60E-16 | 1.37E-14 | 15.7963 |
| Gm33973 | 143.0476 | 10.4904 | 1.2573 | 8.3433 | 7.22E-17 | 6.44E-15 | 16.1414 |
| Grap2 | 122.7195 | 10.4623 | 1.2541 | 8.3422 | 7.29E-17 | 6.47E-15 | 16.1373 |
| Hnf4a | 2846.1873 | 10.4505 | 0.5712 | 18.2972 | 8.71E-75 | 2.00E-71 | 74.0602 |
| Plch1 | 75.7269 | 10.3084 | 1.3214 | 7.8009 | 6.15E-15 | 4.51E-13 | 14.2114 |
| Acot6 | 161.6218 | 10.2949 | 1.2361 | 8.3286 | 8.18E-17 | 7.22E-15 | 16.0873 |
| Top 20 Downregulated | | | | | | | |
| Map1lc3a | 79.8593 | -10.5596 | 1.4301 | -7.3837 | 1.54E-13 | 1.02E-11 | 12.8126 |
| Nrp1 | 86.8761 | -10.6057 | 1.3804 | -7.6832 | 1.55E-14 | 1.11E-12 | 13.8092 |
| Cbr3 | 82.3391 | -10.6238 | 1.3011 | -8.1651 | 3.21E-16 | 2.69E-14 | 15.4933 |
| Man1a | 117.4970 | -10.8553 | 1.2742 | -8.5193 | 1.61E-17 | 1.54E-15 | 16.7944 |
| Slc16a7 | 111.4777 | -10.9863 | 1.2833 | -8.5609 | 1.12E-17 | 1.12E-15 | 16.9510 |
| Ppp1r9a | 144.1616 | -11.2001 | 1.3113 | -8.5411 | 1.33E-17 | 1.31E-15 | 16.8763 |
| Cd200r4 | 165.4777 | -11.6197 | 1.4107 | -8.2367 | 1.77E-16 | 1.51E-14 | 15.7518 |
| Cd86 | 190.6453 | -11.6320 | 1.2592 | -9.2374 | 2.53E-20 | 3.18E-18 | 19.5976 |
| Fam135a | 178.8361 | -11.7375 | 1.2635 | -9.2896 | 1.55E-20 | 2.00E-18 | 19.8100 |
| Ctsf | 961.5254 | -11.8048 | 1.1807 | -9.9978 | 1.56E-23 | 2.53E-21 | 22.8073 |
| Ifi44l | 203.2227 | -11.9245 | 2.9747 | -4.0087 | 6.11E-05 | 0.00105052 | 4.2143 |
| Csf1 | 270.2009 | -12.1645 | 1.3519 | -8.9978 | 2.30E-19 | 2.73E-17 | 18.6378 |
| Ptgs2 | 355.8014 | -12.5397 | 1.4134 | -8.8723 | 7.17E-19 | 7.79E-17 | 18.1447 |
| Ifi44 | 441.1676 | -13.0415 | 2.2671 | -5.7526 | 8.79E-09 | 2.97E-07 | 8.0560 |
| Sdc1 | 588.3062 | -13.2789 | 1.2497 | -10.6260 | 2.26E-26 | 4.41E-24 | 25.6467 |
| Runx3 | 539.4463 | -13.3317 | 1.2124 | -10.9963 | 3.98E-28 | 8.70E-26 | 27.4002 |
| Fads2 | 2937.5049 | -13.9230 | 1.0557 | -13.1889 | 1.02E-39 | 4.67E-37 | 38.9928 |
| Cldn11 | 1221.9780 | -14.1858 | 1.2941 | -10.9618 | 5.83E-28 | 1.25E-25 | 27.2342 |
| Igf1 | 984.2160 | -14.1995 | 1.2138 | -11.6982 | 1.30E-31 | 3.57E-29 | 30.8857 |
| Ly86 | 1421.6730 | -14.7304 | 2.0209 | -7.2889 | 3.13E-13 | 1.97E-11 | 12.5050 |

**Supplemental Table 1. Top 20 upregulated and downregulated genes between cell types.** DESeq2 results of SP-R210$_L$(DN) vs WT cells (a), SP-R210(KO) vs WT (b), and SP-R210(KO) SP-R210$_L$(DN) (c), were sorted by log2foldchange of gene counts; 20 highest and 20 lowest log2foldchanges for each conditions are listed. Additional data can be found in project GEO repository

### Supplemental Table 2

#### a) Upregulated Genes in SP-R210$_L$(DN) cells with PU.1 Associations

| Symbol | baseMean | log2FoldChange | lfcSE | stat | pvalue | padj | log10(pvalue) |
|---|---|---|---|---|---|---|---|
| Associated with Unique WT PU.1 Peaks | | | | | | | |
| 2510009E07Rik | 203.726944 | 11.4879875 | 1.3526278 | 8.493089 | 2.01E-17 | 2.51E-15 | 16.696336 |
| Ly86 | 1421.67295 | 10.8861482 | 1.7479257 | 6.228038 | 4.72E-10 | 2.43E-08 | 9.325771 |
| Runx3 | 539.446275 | 10.0702658 | 0.7789263 | 12.928394 | 3.11E-38 | 1.69E-35 | 37.506888 |
| Pla1a | 161.588704 | 6.9913498 | 0.6996229 | 9.993026 | 1.64E-23 | 3.26E-21 | 22.786449 |
| Maml2 | 28.836878 | 6.252286 | 1.0484955 | 5.963102 | 2.47E-09 | 1.16E-07 | 8.606437 |
| Zbp1 | 32.404636 | 5.221602 | 1.1055733 | 4.722981 | 2.32E-06 | 6.36E-05 | 5.633741 |
| Synpo2 | 17.477322 | 5.199975 | 1.1542397 | 4.505108 | 6.63E-06 | 1.65E-04 | 5.178231 |
| Fads3 | 10.226962 | 4.7217979 | 1.1427345 | 4.132017 | 3.60E-05 | 7.58E-04 | 4.444187 |
| Cxxc5 | 41.271311 | 3.5245589 | 0.7225524 | 4.877929 | 1.07E-06 | 3.14E-05 | 5.969782 |
| Gm13205 | 16.506485 | 2.7732506 | 0.7111879 | 3.899463 | 9.64E-05 | 1.82E-03 | 4.015894 |
| Afap1 | 6.041105 | 2.452814 | 1.1460487 | 2.140235 | 3.23E-02 | 2.25E-01 | 1.490317 |
| Sp110 | 155.070338 | 1.4592475 | 0.5161249 | 2.827315 | 4.69E-03 | 5.11E-02 | 2.328456 |
| Pid1 | 1546.81788 | 1.3704958 | 0.2784811 | 4.921325 | 8.60E-07 | 2.56E-05 | 6.065702 |
| Rgs10 | 1046.38607 | 1.3610551 | 0.3257197 | 4.178608 | 2.93E-05 | 6.32E-04 | 4.53269 |
| Carmil1 | 38.263073 | 1.3609569 | 0.3919021 | 3.472696 | 5.15E-04 | 7.92E-03 | 3.287974 |
| Gpr157 | 49.606685 | 1.3563539 | 0.4921529 | 2.75596 | 5.85E-03 | 6.14E-02 | 2.232695 |
| 1110002J07Rik | 105.214585 | 1.3043878 | 0.28665 | 4.550455 | 5.35E-06 | 1.36E-04 | 5.271403 |
| Fam20c | 4797.09848 | 1.2753051 | 0.462693 | 2.756266 | 5.85E-03 | 6.14E-02 | 2.233101 |
| Ankrd44 | 289.834642 | 1.1859217 | 0.3581261 | 3.311464 | 9.28E-04 | 1.31E-02 | 3.03241 |
| Tmcc3 | 172.526497 | 1.1616585 | 0.4297652 | 2.703007 | 6.87E-03 | 6.99E-02 | 2.162947 |
| Associated with Unique SP-R210L(DN) PU.1 Peaks | | | | | | | |
| Ly86 | 1421.67295 | 10.8861482 | 1.7479257 | 6.228038 | 4.72E-10 | 2.43E-08 | 9.325771 |
| Igf1 | 984.216035 | 10.3632955 | 0.662041 | 15.653556 | 3.14E-55 | 2.68E-52 | 54.502846 |
| Gtpbp10 | 72.173774 | 9.9972327 | 1.2484751 | 8.007555 | 1.17E-15 | 1.19E-13 | 14.931771 |
| Fam135a | 178.836122 | 9.4934333 | 1.1274069 | 8.420592 | 3.75E-17 | 4.45E-15 | 16.426446 |
| Hdac9 | 126.637125 | 9.0052439 | 1.2061133 | 7.466333 | 8.25E-14 | 6.54E-12 | 13.083755 |
| Angptl2 | 974.492245 | 8.8213905 | 0.6868856 | 12.842591 | 9.47E-38 | 4.71E-35 | 37.023867 |
| Unc5b | 31.5345 | 8.798551 | 1.3956567 | 6.304237 | 2.90E-10 | 1.53E-08 | 9.538177 |
| Cacna1d | 29.671987 | 8.7022819 | 1.2910069 | 6.740693 | 1.58E-11 | 9.71E-10 | 10.802354 |
| Csf3r | 25.419805 | 8.4844854 | 1.3931977 | 6.089937 | 1.13E-09 | 5.61E-08 | 8.947093 |
| Pde8a | 321.259807 | 8.2865551 | 0.6484095 | 12.779817 | 2.13E-37 | 1.00E-34 | 36.672498 |
| Zfp503 | 16.945437 | 7.9112174 | 1.5159621 | 5.218612 | 1.80E-07 | 6.24E-06 | 6.744078 |
| Rtp4 | 270.719967 | 7.7392012 | 2.2202541 | 3.485728 | 4.91E-04 | 7.58E-03 | 3.309095 |
| Vat1l | 14.697583 | 7.7060883 | 1.7039346 | 4.522526 | 6.11E-06 | 1.54E-04 | 5.213916 |
| AI504432 | 13.428983 | 7.5663189 | 1.4125576 | 5.356467 | 8.49E-08 | 3.11E-06 | 7.071273 |
| Tnip3 | 320.224052 | 7.166918 | 0.7427586 | 9.649055 | 4.96E-22 | 8.63E-20 | 21.304422 |
| Dab2ip | 67.111512 | 7.1070825 | 0.9160959 | 7.758011 | 8.63E-15 | 7.66E-13 | 14.064133 |
| Tent5c | 92.311213 | 7.0367245 | 1.0276478 | 6.847408 | 7.52E-12 | 4.81E-10 | 11.123783 |
| Slc22a17 | 69.199366 | 6.3582987 | 0.8700843 | 7.307681 | 2.72E-13 | 2.04E-11 | 12.565763 |
| Scin | 5.758523 | 6.3446806 | 2.6134111 | 2.427739 | 1.52E-02 | 1.28E-01 | 1.818349 |
| Maml2 | 28.836878 | 6.252286 | 1.0484955 | 5.963102 | 2.47E-09 | 1.16E-07 | 8.606437 |

#### b) Downregulated Genes in SP-R210$_L$(DN) cells with PU.1 Associations

| Symbol | baseMean | log2FoldChange | lfcSE | stat | pvalue | padj | log10(pvalue) |
|---|---|---|---|---|---|---|---|
| Associated with Unique WT PU.1 Peaks | | | | | | | |
| Eif2s3y | 7.08E+02 | -12.346023 | 1.2404609 | -9.952771 | 2.45E-23 | 4.83E-21 | 22.610376 |
| Rasd2 | 2.86E+02 | -12.1703 | 1.3372706 | -9.100851 | 8.96E-20 | 1.40E-17 | 19.047561 |
| Marcks | 2.50E+02 | -11.69022 | 1.3766542 | -8.491762 | 2.04E-17 | 2.52E-15 | 16.691378 |
| Rnf217 | 2.15E+02 | -10.521487 | 1.3372855 | -7.867794 | 3.61E-15 | 3.37E-13 | 14.442552 |
| Hnf4a | 2.85E+03 | -10.223966 | 0.5712191 | -17.898501 | 1.21E-71 | 2.71E-68 | 70.916714 |
| Tmc5 | 1.40E+02 | -10.216049 | 1.2167494 | -8.396181 | 4.61E-17 | 5.44E-15 | 16.33608 |
| Prss22 | 1.00E+02 | -9.2884558 | 1.2250052 | -7.582381 | 3.39E-14 | 2.84E-12 | 13.469454 |
| Alpk3 | 1.68E+02 | -9.2756254 | 1.2018314 | -7.717909 | 1.18E-14 | 1.03E-12 | 13.927185 |
| Vopp1 | 1.23E+02 | -9.1739127 | 1.0807465 | -8.488497 | 2.09E-17 | 2.57E-15 | 16.679174 |
| Colq | 6.40E+01 | -9.0734916 | 1.2794878 | -7.091503 | 1.33E-12 | 9.11E-11 | 11.87725 |
| Ptprr | 5.87E+01 | -9.0247646 | 1.266043 | -7.128324 | 1.02E-12 | 7.09E-11 | 11.993113 |
| Myzap | 3.01E+02 | -8.5154328 | 0.7855288 | -10.840383 | 2.22E-27 | 6.11E-25 | 26.654544 |
| Vil1 | 8.06E+02 | -8.511351 | 0.5215779 | -16.318465 | 7.29E-60 | 8.57E-57 | 59.136992 |
| Fgf6 | 9.20E+01 | -8.4900598 | 1.261077 | -6.732388 | 1.67E-11 | 1.02E-09 | 10.777542 |
| Ccl6 | 1.60E+04 | -8.4864471 | 0.3281609 | -25.860628 | 1.85E-147 | 1.66E-143 | 146.733355 |
| Cntnap5c | 2.53E+02 | -8.4491439 | 1.0806995 | -7.818217 | 5.36E-15 | 4.83E-13 | 14.271025 |
| Cgnl1 | 1.59E+02 | -8.3811778 | 0.8691499 | -9.64296 | 5.26E-22 | 9.08E-20 | 21.278624 |
| Plvap | 5.67E+02 | -8.3680443 | 0.6797555 | -12.310374 | 7.97E-35 | 3.04E-32 | 34.098797 |
| Bicd1 | 4.71E+01 | -8.1356791 | 1.2479781 | -6.519088 | 7.07E-11 | 3.95E-09 | 10.150358 |
| Espn | 2.42E+01 | -7.3760433 | 1.5182786 | -4.858162 | 1.18E-06 | 3.44E-05 | 5.926353 |
| Associated with Unique SP-R210L(DN) Peaks | | | | | | | |
| Marcks | 250.241782 | -11.69022 | 1.3766542 | -8.491762 | 2.04E-17 | 2.52E-15 | 16.691378 |
| Rgs9 | 76.293252 | -9.579173 | 1.2203352 | -7.849624 | 4.17E-15 | 3.84E-13 | 14.379567 |
| 1700016C15Rik | 2591.7708 | -9.5013025 | 0.6912432 | -13.745239 | 5.44E-43 | 3.75E-40 | 42.264456 |
| Hsd17b2 | 618.52305 | -8.8367303 | 0.8188342 | -10.791843 | 3.76E-27 | 1.02E-24 | 26.424617 |
| Lyz1 | 97226.6007 | -8.6093753 | 0.4260457 | -20.207633 | 8.39E-91 | 2.51E-87 | 90.076371 |
| Vil1 | 805.646244 | -8.511351 | 0.5215779 | -16.318465 | 7.29E-60 | 8.57E-57 | 59.136992 |
| Plekha6 | 486.69065 | -8.1476721 | 1.2629229 | -6.451441 | 1.11E-10 | 6.06E-09 | 9.955493 |
| Fa2h | 57.021779 | -7.3643767 | 1.1027277 | -6.678328 | 2.42E-11 | 1.45E-09 | 10.616752 |
| Fgr | 571.714127 | -7.0323223 | 0.5181006 | -13.573276 | 5.77E-42 | 3.69E-39 | 41.238936 |
| Fam49a | 5.356929 | -6.4981335 | 1.9375865 | -3.353726 | 7.97E-04 | 1.15E-02 | 3.098371 |
| 1810046K07Rik | 4.715197 | -6.310071 | 1.8062176 | -3.493528 | 4.77E-04 | 7.41E-03 | 3.32177 |
| Rhpn2 | 685.880148 | -6.3078935 | 0.3629226 | -17.380824 | 1.15E-67 | 1.72E-64 | 66.938225 |
| Padi1 | 7.865312 | -6.2755914 | 1.5731342 | -3.989228 | 6.63E-05 | 1.31E-03 | 4.178561 |
| Stc1 | 4.245222 | -6.1584502 | 2.7647695 | -2.227473 | 2.59E-02 | 1.91E-01 | 1.586438 |
| Pbx4 | 7.593239 | -6.1019118 | 1.5277681 | -3.994004 | 6.50E-05 | 1.29E-03 | 4.187309 |
| Rasgrf2 | 3.733174 | -5.8983628 | 1.814863 | -3.250032 | 1.15E-03 | 1.58E-02 | 2.937824 |
| Zc3h12d | 224.320895 | -5.826769 | 0.4408165 | -13.218128 | 6.90E-40 | 4.26E-37 | 39.161408 |
| Tprg | 5.325286 | -5.6881678 | 1.465125 | -3.882377 | 1.03E-04 | 1.94E-03 | 3.98531 |
| Trib2 | 5.300092 | -5.6096201 | 1.953552 | -2.871498 | 4.09E-03 | 4.57E-02 | 2.388774 |
| Lmntd1 | 4.018488 | -5.5473629 | 2.2762666 | -2.437044 | 1.48E-02 | 1.26E-01 | 1.829508 |

**Supplemental Table 2. Top 20 upregulated and downregulated genes with associations with unique WT or SP-R210$_L$(DN) PU.1 peaks.** DESeq2 results of SP-R210$_L$(DN) vs WT cells were sorted by log2foldchange of gene counts. Upregulated (a) and downregulated (2) genes were filtered by genes that had PU.1 binding regions that had PU.1 peaks unique to either WT or SP-R210$_L$(DN) cells. 20 Highest and 20 lowest log2foldchanges for each conditions are listed. Additional data can be found in project GEO repository
